# Single cell and single nucleus RNA-Seq reveal cellular heterogeneity and homeostatic regulatory networks in adult mouse stria vascularis

**DOI:** 10.1101/756635

**Authors:** Soumya Korrapati, Ian Taukulis, Rafal Olszewski, Madeline Pyle, Shoujun Gu, Riya Singh, Carla Griffiths, Daniel Martin Izquierdo, Erich Boger, Robert J. Morell, Michael Hoa

## Abstract

The stria vascularis (SV) generates the endocochlear potential (EP) in the inner ear and is necessary for proper hair cell mechanotransduction and hearing. While channels belonging to SV cell types are known to play crucial roles in EP generation, relatively little is known about gene regulatory networks that underlie the ability of the SV to generate and maintain the EP. Using single cell and single nucleus RNA-sequencing, we identify and validate known and rare cell populations in the SV. Furthermore, we establish a basis for understanding molecular mechanisms underlying SV function by identifying potential gene regulatory networks as well as druggable gene targets. Finally, we associate known deafness genes with adult SV cell types. This work establishes a basis for dissecting the genetic mechanisms underlying the role of the SV in hearing and will serve as a basis for designing therapeutic approaches to hearing loss related to SV dysfunction.

## Introduction

Ionic homeostasis in the endolymph-containing compartment of the cochlea, the scala media, is a critical factor in enabling proper hair cell mechanotransduction and hearing (Hibino, Nin, Tsuzuki, & Kurachi, 2010; Wangemann, 2002, 2006). The endolymph is the atypical potassium rich extracellular fluid of the cochlear duct. This high potassium concentration results in a +80 millivolt (mV) positive potential known as the endocochlear potential (EP) (Patuzzi, 2011; Wangemann, 2002). The stria vascularis (SV), a non-sensory epithelial tissue in the lateral wall of the cochlea, generates and maintains this high potassium concentration and the EP.

The SV is a complex, heterogenous tissue consisting of several cell types that work together to generate and maintain the EP. Cell types identified as critical to this role thus far include marginal cells, intermediate cells and basal cells (Gow, 2004; Marcus, Wu, Wangemann, & Kofuji, 2013; Wangemann, 2002; Wangemann et al., 2004). The marginal cells (MCs) face the endolymph and extend basolateral projections that interdigitate with the intermediate cells (ICs) which have projections that run in both directions towards marginal cells apically and basal cells at the basolateral end (Figure 1A). Basal cells (BCs) are connected to each other by tight junctions (like the MCs) to prevent leakage of ions (Kitajiri et al., 2004). At least two of these cell types, marginal and intermediate cells appear to have densely interdigitating processes (Figure 1B) intimating at the close functional interaction between these cell types in the SV (Nakazawa, Spicer, & Schulte, 1995; Steel & Barkway, 1989). In addition, other cell types in the stria vascularis include spindle cells, macrophages, pericytes and endothelial cells (Taku Ito et al., 2014; Neng, Zhang, Kachelmeier, & Shi, 2013; Shi, 2016).

**Figure 1.**
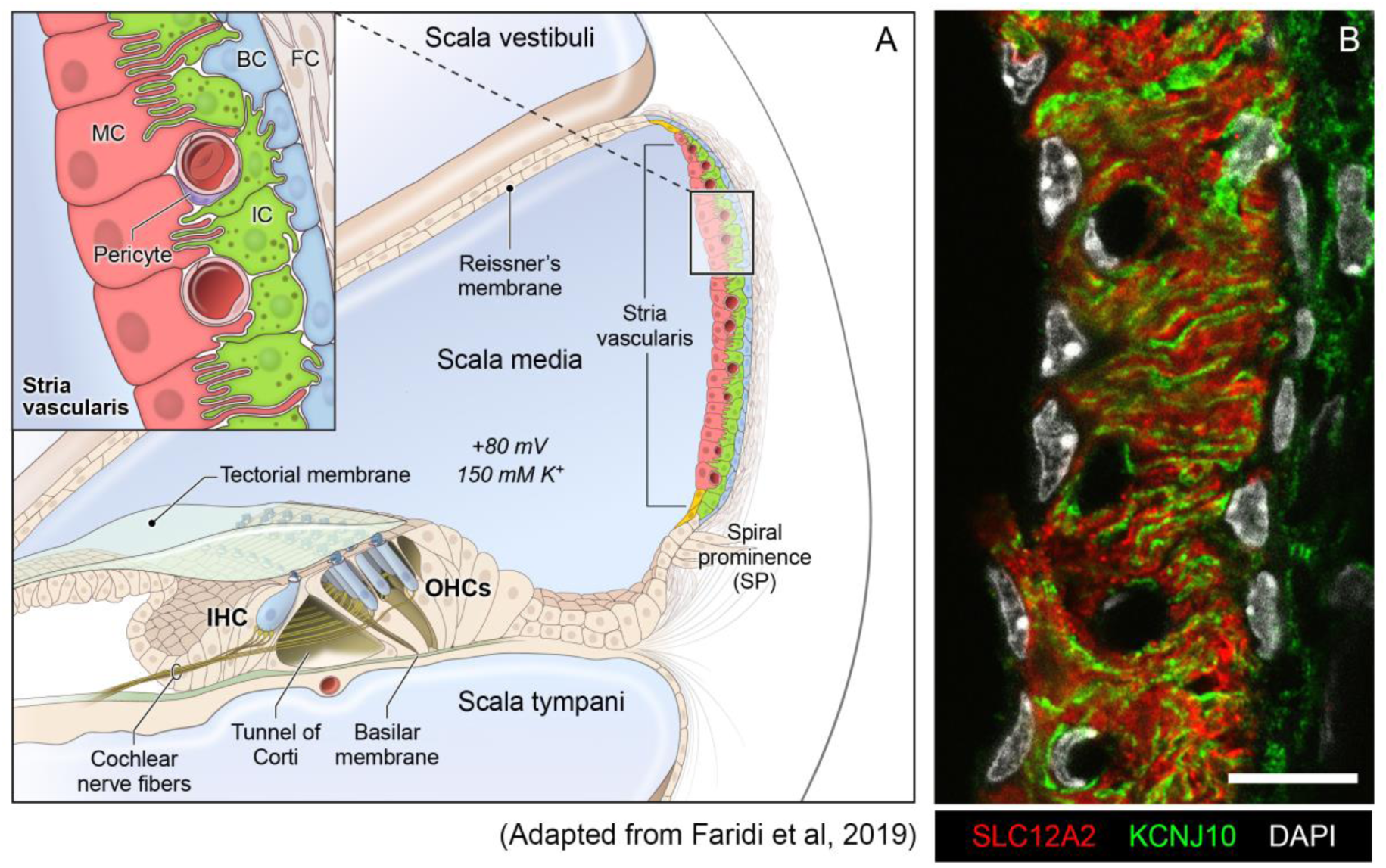
Stria vascularis cellular heterogeneity and organization. (A), Schematic of the stria vascularis (SV) and its relationship to structures in the cochlea. The SV is composed of three layers of cells and is responsible for generating the +80 mV endocochlear potential (EP) and the high potassium concentration in the endolymph-containing scala media. The relationship between the marginal, intermediate and basal cells are demonstrated with the marginal extending basolateral projections to interdigitate with intermediate cells, which have bidirectional cellular projections that interdigitate with both marginal and basal cells. In addition to these cell types, other cell types, including spindle cells (yellow), endothelial cells, pericytes, and macrophages (not shown) are present in the SV. **(B),** Cross-section of the SV in a postnatal day 30 (P30) mouse immunostained with anti-SLC12A2 (marginal cells, red), anti-KCNJ10 (intermediate cells, green), and DAPI (4′,6-diamidino-2-phenylindole) for nuclei. Notice the interdigitation of cellular processes from both intermediate and marginal cells. Scale bar is 20 μm.

Knowledge regarding the role of the three main cell types (MCs, ICs, BCs) in the generation and maintenance of the EP is based on previous work by others. Mutations in genes expressed by marginal, intermediate and basal cells in the SV are known to cause deafness and dysfunction in EP generation. In marginal cells, mutations in Kcnq1, Kcne1 and Barttin (Bsnd) result in a loss or reduction of EP and deafness (Chang et al., 2015; Faridi et al., 2019; Rickheit et al., 2008). *Kcnq1/Kcne1* encode the voltage-gated potassium channel Kv7.1 and play a crucial role in secreting potassium and maintaining the EP. Conditional *Kcnq1* null mice exhibit collapsed Reissner’s membrane, loss of EP, and are deaf (Chang et al., 2015). Barttin (*Bsnd*) is a beta subunit of chloride channel ClC-K, mutations in which cause deafness and Bartter syndrome IV in humans. Conditional null mice of barttin in the inner ear exhibit hearing loss with reduced EP (Riazuddin et al., 2009; Rickheit et al., 2008). In intermediate cells, *Kcnj10* encodes Kir4.1, an inwardly rectifying potassium channel, which is necessary for the generation of the EP. Loss or mutations in *Kcnj10* have been shown to cause hearing loss in humans and mice, accompanied by an absence of EP and loss of endolymphatic potassium (J. Chen & Zhao, 2014; Marcus et al., 2013; Wangemann et al., 2004). Finally, basal cells play a role in barrier formation and prevent ion leakage from the SV. Claudin 11 (*Cldn11*), a tight junction protein expressed in SV basal cells, is critical to this function as demonstrated by deafness and low EP in *Cldn11* null mice (Gow, 2004; Kitajiri et al., 2004).

Despite continuing interest in SV cell types, an understanding of cellular heterogeneity, including a comprehensive understanding of SV cell type-specific transcriptional profiles, is incomplete. Furthermore, the mechanisms by which the various cell types contribute to EP generation as well as other strial functions remains largely undefined (Ohlemiller, 2009). Recently, both single cell and single nucleus approaches have been utilized to define transcriptional profiles of cells from organs and tissues with significant cellular heterogeneity (Wu, Kirita, Donnelly, & Humphreys, 2019; Zeng et al., 2016). Given the presence of a heterogeneous group of cell types with significant cell size and shape heterogeneity, we set out to define the transcriptional profiles of the three major cell types implicated in EP generation by utilizing single cell RNA-Seq (scRNA-Seq) and single nucleus RNA-Seq (snRNA-Seq) in the adult SV. In doing so, we seek to define transcriptional heterogeneity between SV cell types and define gene regulatory networks in the unperturbed wild type adult SV that can serve as a basis for investigating mechanisms responsible for SV functions.

## Results

### Defining cellular heterogeneity within the adult stria vascularis (SV)

The SV is composed of a heterogeneous group of cell types that work together to generate the endocochlear potential. Defining these cell types and their respective transcriptional profiles in the adult mammalian SV is a critical first step towards understanding the genetic mechanisms that produce the EP. Cell isolation for single cell and single nucleus approaches must be optimized for the tissues they are targeting. In particular, cellular heterogeneity within a tissue can be manifested, not just by cell type heterogeneity, but also heterogeneity in size and shape, potentially necessitating different techniques to gain a comprehensive perspective on a given tissue. As can be seen in the representative image of the SV in Figure 1B, marginal cells (red), with cell nuclei eccentrically located closer to the endolymph (apical-medial), interdigitate with intermediate cells (green) with cell nuclei arranged so that processes extend to both the apical-medial and basolateral surfaces of the SV. The basal cells, which line the basolateral surface of the stria vascularis, appear to be relatively flat cells. The marginal, basal, and spindle cells appear to have relatively flat-appearing nuclei, while those of the intermediate cells appear more oblong in shape. For the stria vascularis, in order to address the challenge of transcriptional profiling a tissue composed of heterogeneous cell types with significant heterogeneity in cell size and shape, we utilized two methods of transcriptional profiling, single cell RNA-Seq (scRNA-Seq) and single nucleus RNA-Seq (snRNA-Seq).

#### Stria vascularis (SV) cell types exhibit clear transcriptional differences

Unbiased clustering of adult SV single cells and nuclei was performed independently. After unbiased clustering of cells by shared gene expression, known cell type-specific markers were utilized to identify these agnostically determined cell clusters. Based on these markers, marginal cell, intermediate cell, basal cell, spindle/root cell, macrophages, and immune and hematopoietic cell clusters were identified within the adult SV scRNA-Seq and snRNA-Seq datasets (Figure 2A). Here, we will focus the analyses on the marginal, intermediate, basal, and spindle/root cell clusters. Feature plots of both scRNA-Seq and snRNA-Seq datasets demonstrate the correlation of known gene expression for marginal cells (*Kcne1*, *Kcnq1*) (Wangemann, 2002), intermediate cells (*Cd44*, *Met*) (Rohacek et al., 2017; Shibata, Miwa, Wu, Levitt, & Ohyama, 2016), basal cells (*Cldn11*, *Tjp1*) (Gow, 2004; Lee, Shin, Sagong, Kim, & Bok, 2017; W. Liu, Schrott-Fischer, Glueckert, Benav, & Rask-Andersen, 2017), and spindle/root cells (*Slc26a4*) between the two datasets (Figure 2B, 2D) (Nishio et al., 2016). Violin plots demonstrate relative expression of these known SV cell type-specific genes across the 4 main cell types (Figure 2C, 2E). Confirmatory smFISH and immunohistochemistry demonstrates expression of these known markers for marginal cells (*Kcne1*, *Kcnq1*), intermediate cells (*Cd44*, *Met*), basal cells (CLDN11, ZO-1), and spindle/root cells (*Slc26a4*) (Figure 1F). Co-expression of *Kcne1* and *Kcnq1* RNA to marginal cells of the adult SV can be seen, particularly in close proximity to marginal cell nuclei (Figure 2F). *Cd44* and *Met* RNA is co-expressed in intermediate cell nuclei (Figure 2F). CLDN11 and ZO-1 (the protein product of the *Tjp1* gene) are co-expressed in basal cells (Figure 2F). The confirmation of cell type clusters with known gene expression across both datasets strengthens the validity of the unbiased clustering as well as the capability of both approaches to assess transcriptome profiles in the SV.

**Figure 2.**
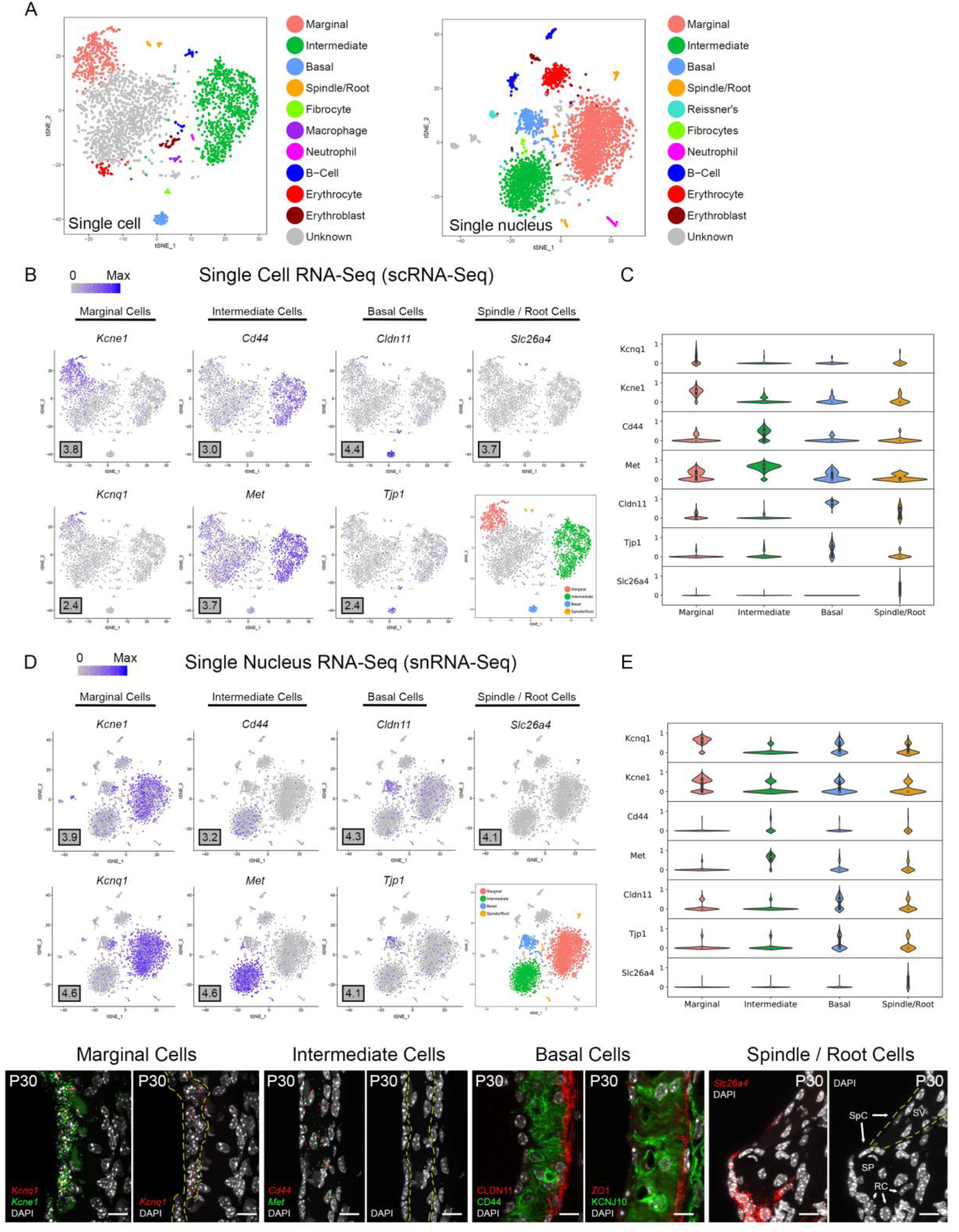
Stria vascularis (SV) cell types show clear transcriptional differences at the single cell and single nucleus level. **(A)**, Unbiased clustering was performed on both single cell RNA-Seq (scRNA-Seq) and single nucleus RNA-Seq (snRNA-Seq) datasets from the P30 mouse SV. Known cell type-specific markers were utilized to identify these agnostically determined cell clusters. Based on these known markers, clusters consisting of marginal cells, intermediate cells, basal cells, spindle / root cells, and immune cell types, including macrophages, B cells, and neutrophils, were identified. tSNE plot of P30 mouse SV scRNA-Seq dataset demonstrates clustering of SV cell types (LEFT panel). tSNE plot of P30 mouse SV snRNA-Seq dataset demonstrates clustering of SV cell types with similar identification of cell type-specific clusters (RIGHT panel). **(B),** Feature plots for P30 SV scRNA-Seq data demonstrate expression of known markers for 4 main SV cell types: marginal cells (*Kcne1*, *Kcnq1*), intermediate cells (*Cd44*, *Met*), basal cells (*Cldn11*, *Tjp1*), and spindle / root cells (*Slc26a4*). 4 main cell types are highlighted on the inset tSNE plot of the scRNA-Seq dataset. **(C),** Grid violin plot of scRNA-Seq dataset demonstrates expression of known marker genes amongst SV cell types. Normalized counts were scaled to a range of 0 to 1 using min-max scaling. **(D),** Feature plots for P30 SV snRNA-Seq data demonstrate expression of known markers for 4 main SV cell types: marginal cells (*Kcne1*, *Kcnq1*), intermediate cells (*Cd44*, *Met*), basal cells (*Cldn11*, *Tjp1*), and spindle / root cells (*Slc26a4*). 4 main cell types are highlighted on the inset tSNE plot of the snRNA-Seq dataset. **(E),** Grid violin plot of snRNA-Seq dataset demonstrates expression of known marker genes amongst SV cell types. Normalized counts were scaled to a range of 0 to 1 using min-max scaling. **(F),** Validation of known gene expression was performed using single molecule fluorescent *in situ* hybridization (smFISH) and fluorescence immunohistochemistry. Known marginal cell-specific transcripts (*Kcne1*, *Kcnq1*) and intermediate cell-specific transcripts (*Cd44*, *Met*) are co-localized by smFISH. Yellow dashed lines outline the SV marginal cell layer showing Kcnq1 transcript expression in marginal cells with DAPI to label nuclei. Yellow dashed lines outline the SV intermediate cell layer in the image with DAPI only. CLDN11 and ZO-1 (protein product of *Tjp1*) are localized to the basal cell layer with anti-CD44 and anti-KCNJ10 immunostaining to label the adjacent intermediate cell layer for contrast, respectively. Finally, *Slc26a4* transcripts are localized to the spindle / root cells by smFISH. Yellow dashed lines in the image to the right with only the DAPI labeling demarcate the SV and spindle cells (SpC) and root cells (RC) in the spiral prominence (SP) are identified. DAPI (4′,6-diamidino-2-phenylindole). All scale bars 10 μm.

#### Novel cell type-specific genes are identified by scRNA-Seq and snRNA-Seq of the adult stria vascularis (SV)

Based on the transcriptional profiles, novel cell type-specific genes were identified and validated in adult SV. Feature plots demonstrate expression of selected novel candidate genes in marginal cells (*Abcg1*, *Heyl*), intermediate cells (*Nrp2*, *Kcnj13*), basal cells (*Sox8*, *Nr2f2*), and spindle/root cells (*P2rx2*, *Kcnj16*) in both scRNA-Seq and snRNA-Seq datasets, respectively (Figure 3A, 3C). Violin plots demonstrate the relative expression of these novel SV cell type-specific genes across the 4 main cell types (Figure 3B, 3D). The violin plots highlight aspects of shared gene expression between marginal cells and spindle cells amongst the selected candidate genes. This trend of shared gene expression was observed between marginal and spindle cells in terms of shared groups of genes (Supplemental Figure S1). Representative images depict co-expression of candidate gene RNA by smFISH in marginal cells (*Abcg1*, *Heyl*), intermediate cells (*Nrp2*, *Kcnj13*), basal cells (*Sox8*, *Nr2f2*) and spindle / root cells (*P2rx2*, *Kcnj16*) (Figure 3E, 3F). None of these gene transcripts have been previously identified in the mammalian SV. A discussion of the novel genes identified in this study is provided (Supplemental Results).

**Figure 3.**
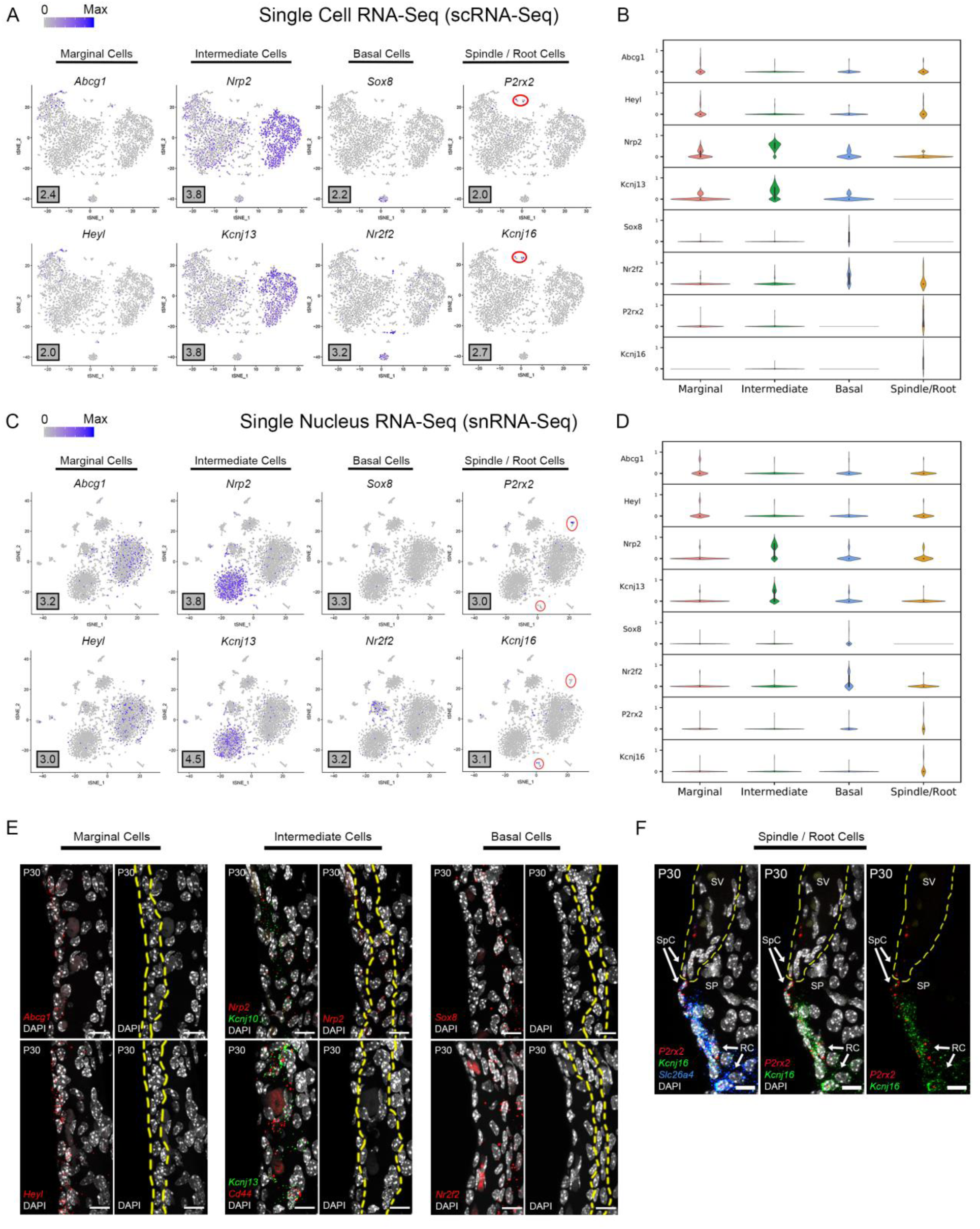
Single cell and single nucleus transcriptome profiles reveal common novel cell type-specific genes in the adult stria vascularis. **(A)**, Feature plots for P30 SV scRNA-Seq data demonstrate expression of novel markers for 4 main SV cell types: marginal cells (*Abcg1, Heyl*), intermediate cells (*Nrp2, Kcnj13*), basal cells (*Sox8, Nr2f2*), and spindle / root cells (*P2rx2, Kcnj16*). **(B),** Grid violin plot of scRNA-Seq dataset demonstrates expression of novel marker genes amongst SV cell types. Normalized counts were scaled to a range of 0 to 1 using min-max scaling. **(C),** Feature plots for P30 SV snRNA-Seq data demonstrate expression of novel markers for 4 main SV cell types: marginal cells (*Abcg1, Heyl*), intermediate cells (*Nrp2, Kcnj13*), basal cells (*Sox8, Nr2f2*), and spindle / root cells (*P2rx2, Kcnj16*). **(D),** Grid violin plot of snRNA-Seq dataset demonstrates expression of novel marker genes amongst SV cell types. Normalized counts were scaled to a range of 0 to 1 using min-max scaling. **(E),** Validation of known gene expression was performed using single molecule fluorescent *in situ* hybridization (smFISH). Novel marginal cell-specific transcript (*Abcg1*, *Heyl*) expression in red is demonstrated in and around the marginal cell nuclei on the apical surface of the marginal cells. Novel intermediate cell-specific transcript (*Nrp2*, *Kcnj13*) expression in red is co-expressed with *Kcnj10* and *Cd44* transcripts in green, respectively. Novel basal cell-specific transcript (*Sox8*, *Nr2f2*) expression in red is localized to the basal cell layer. Yellow dashed lines in the images with only the DAPI labeling for nuclei demarcate the respective cell layers (marginal, intermediate and basal cell layers). **(F),** Novel spindle / root cell-specific transcripts, *P2rx2* (in red) and *Kcnj16* (in green) are co-expressed with *Slc26a4* transcripts to spindle and root cells (left panel). The same image without the Slc26a4 (blue) channel (center panel) and without DAPI (right panel) are shown. Note the slightly different distributions in *P2rx2* (red) versus Kcnj16 (green) transcripts amongst spindle and root cells, respectively. Yellow dashed lines demarcate the SV, spindle cells (SpC) and root cells (RC) in the spiral prominence (SP). DAPI (4′,6-diamidino-2-phenylindole). All scale bars 10 μm.

### Defining potential gene regulatory networks in SV marginal, intermediate and basal cells

Focusing on strial marginal, intermediate and basal cells, we sought to identify cell type-specific gene regulatory networks that might serve as a basis for investigating mechanisms related to strial function and eventually on strial dysfunction in human disorders. To identify these potential gene regulatory networks in the adult mouse, two methods (WGCNA, SCENIC) of unbiased gene regulatory network identification were utilized as described previously. WGCNA identifies modules of co-expressed genes without making a link between transcription factors and co-expressed genes. In doing so, WGCNA casts a wider net as it relates to co-expression analysis, potentially identifying indirectly linked genes in a regulatory network. Conversely, SCENIC identifies regulons composed of co-expressed transcription factors and their downstream targets as determined by motif enrichment. This approach identifies transcription factors with potentially directly linked genes in a regulatory network.

#### WGCNA identifies cell type-specific gene regulatory networks in the adult stria vascularis

Topological overlap (TOM) plots for scRNA-Seq and snRNA-Seq datasets from the adult stria vascularis demonstrate clustering of gene modules identified in WGCNA (Figure 4A). The more red a box is, the higher the Pearson correlation coefficient is between modules. The adjacency plots take this adjacency matrix and display highly correlated gene modules (Figure 4B). Cell type-specific modules for marginal, intermediate and basal cells were reprojected onto the feature plots for each dataset (Figure 4C). The top gene ontology (GO) biological process, cellular component and molecular function are summarized in Supplemental Figure S2.

**Figure 4.**
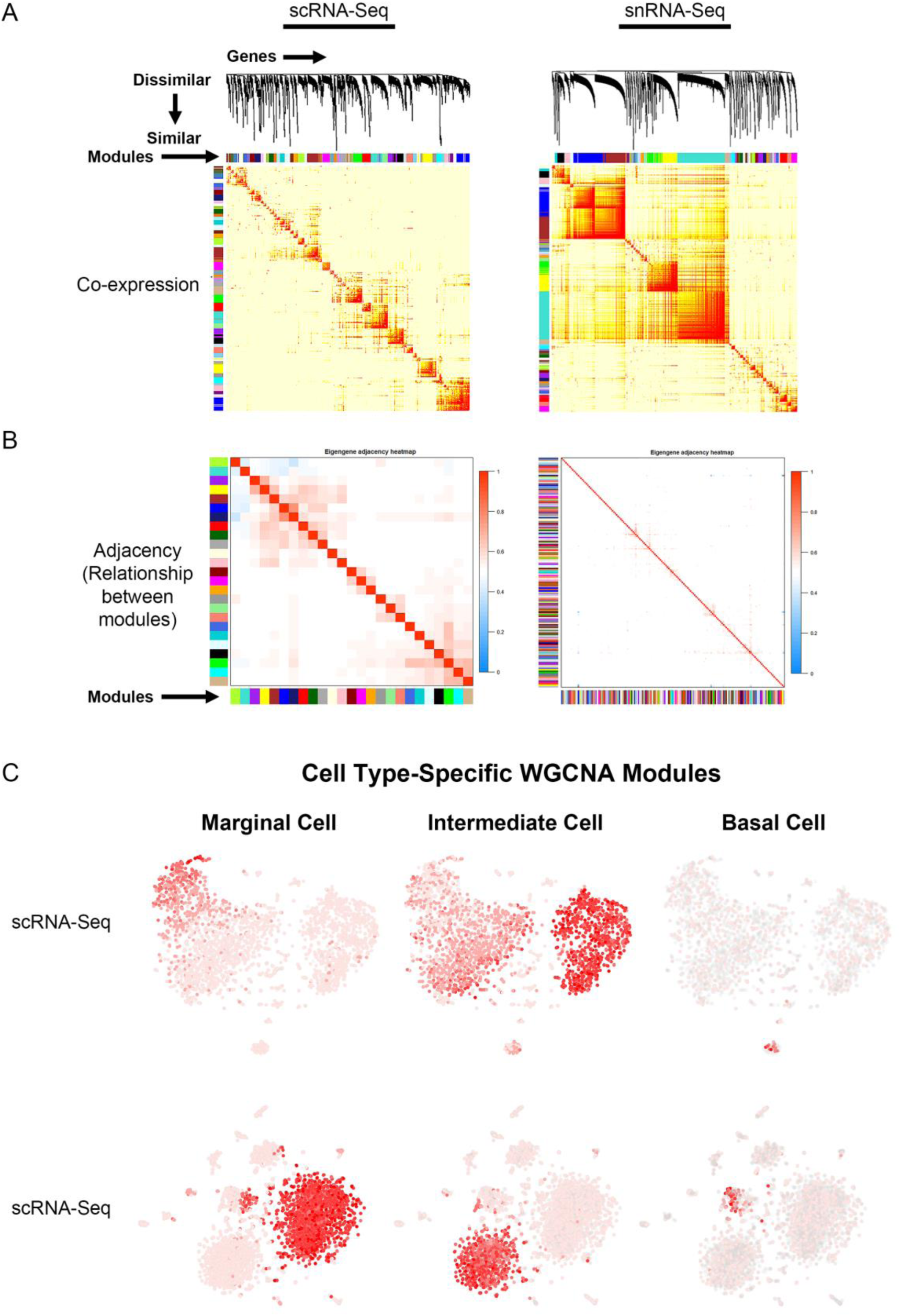
Gene regulatory network inference using weighted gene co-expression network analysis (WGCNA) reveals cell type-specific modules. **(A)**, Topological overlay map (TOM) plots demonstrate clustering of potential gene regulatory networks consisting of co-expressed genes termed modules arrayed along the horizontal and vertical axis. The more red a box is, the higher the Pearson correlation coefficient is between the respective modules. TOM plots are shown for the scRNA-Seq (LEFT) and snRNA-Seq (RIGHT) datasets. **(B),** Adjacency plot displays highly correlated modules in the scRNA-Seq dataset (LEFT panel). Note that in the snRNA-Seq dataset (RIGHT panel), the correlation between modules is minimal. Using known cell type-specific marker genes, we focused on the cell type-specific modules identified in the scRNA-Seq dataset. (C), Cell type-specific WGCNA modules were then projected onto the scRNA-Seq feature plot to demonstrate aggregate cell type-specificity of these WGCNA modules for the three major cell types (marginal, intermediate, and basal cells). The more red a dot is in the feature plot, the greater the aggregate expression of the cell type-specific modules in that given cell.

GO biological process analysis revealed a significant enrichment for genes involved in positive regulation of potassium ion transport (GO:1903288) in SV marginal cells and regulation of potassium ion transport by positive regulation of transcription from RNA polymerase II promoter (GO:0097301) for SV intermediate cells. Additionally, GO biological process analysis revealed a significant enrichment in genes involved in protein stabilization (GO:0050821), neutrophil degranulation (GO:0043312), and positive regulation of rhodopsin gene expression (GO:0045872) in SV marginal, intermediate, and basal cells, respectively. GO molecular function analysis revealed a significant enrichment for genes involved in calcium-transporting ATPase activity (GO:0005388) and translation factor activity, RNA binding (GO:0008135) in SV marginal and intermediate cells, respectively. GO molecular function analysis did not reveal a significantly enriched process in SV basal cells. GO cellular component analysis revealed a significant enrichment for genes involved in G-protein coupled receptor complex (GO:0097648), interleukin-28 receptor complex (GO:0032002), and platelet alpha granule lumen (GO:0031093) for SV marginal, intermediate, and basal cells, respectively. Overall, these analyses affirm the importance of ion homeostasis amongst marginal and intermediate cells and suggest functions related to the immune system (neutrophil degranulation, interleukin-28 receptor complex).

#### SCENIC identifies cell type-specific gene regulatory networks in the adult stria vascularis

TSNE plots demonstrate a comparison between principal component analysis (PCA)-based clustering within Seurat and regulon-based clustering within SCENIC for both scRNA-Seq in the upper panel (Figure 5A) and snRNA-Seq in the lower panel (Figure 5B). Regulon-based clustering within SCENIC identifies similar clusters to those seen within Seurat based on PCA. Display of regulon activity matrix for both scRNA-Seq and snRNA-Seq datasets demonstrates cell type-specific regulons prominently within marginal, intermediate and basal cells in pink, green and blue, respectively. The top gene ontology (GO) biological process, cellular component and molecular function are summarized in Supplemental Figure S2.

**Figure 5.**
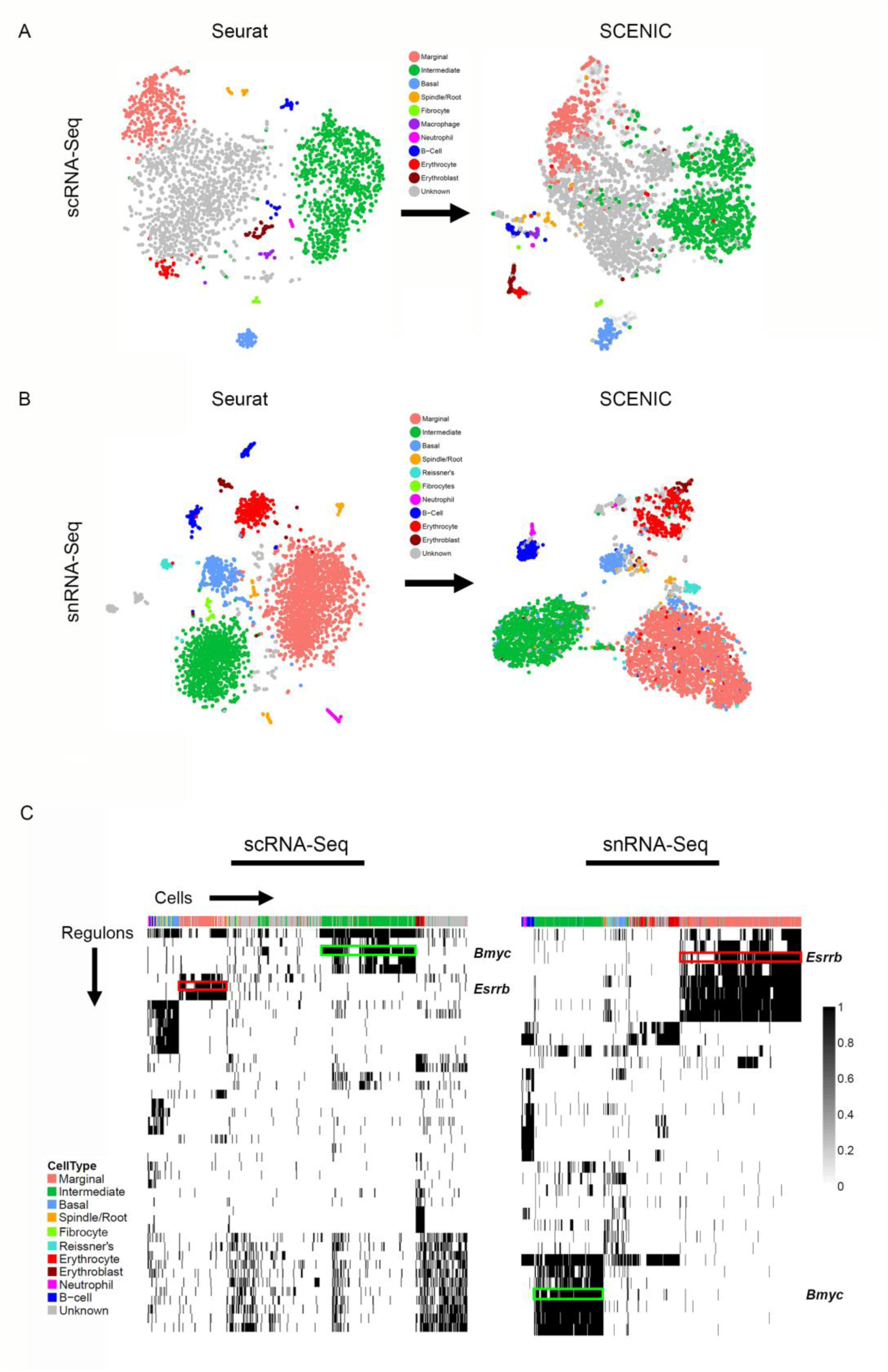
Gene regulatory network inference with SCENIC reveals cell type-specific SCENIC regulons in both scRNA-Seq and snRNA-Seq datasets. **(A)**, Unbiased clustering in Seurat (left panel) compares similarly to regulon-based clustering in SCENIC (right panel) for the scRNA-Seq dataset, resulting in the identification of the same cell type-specific clusters. **(B),** Unbiased clustering in Seurat (left panel) compares similarly to regulon-based clustering in SCENIC (right panel) for the snRNA-Seq dataset, resulting in the identification of the same cell type-specific clusters. **(C),** Regulon heatmaps display cell type-specific regulons identified from both the scRNA-Seq (left panel) and snRNA-Seq (right panel) datasets. *Esrrb* (red box) and *Bmyc* (green box) regulons are demarcated in both datasets (regulon labels to right of heatmap. Histogram demonstrating normalized regulon activity score (scaled from 0 to 1) is shown to right of both heatmaps.

GO biological process analysis for all SV cells reveals a significant enrichment for genes involved in the regulation of transcription from RNA polymerase II promoter (GO:0006357). GO molecular function analysis revealed a significant enrichment for genes involved in protein kinase activity (GO:0019901) and calcium channel regulator activity (GO:0005246) in SV marginal cells, for genes involved in RNA polymerase II regulatory region sequence-specific DNA binding (GO:0000977) and proton-transporting ATPase activity, rotational mechanism (GO:0046961) in SV intermediate cells, and for genes involved in RNA binding (GO:0003723) and phosphatidylinositol phosphate kinase activity (GO:0016307) in SV basal cells. GO cellular component analysis reveals a significant enrichment for genes related to mitochondrion (GO:0005739) and ruffle membrane (GO:0032587) in marginal cells, lysosome (GO:0005764) and late endosome (GO:0005770) in intermediate cells, and focal adhesion (GO:0005925) and actin cytoskeleton (GO:0015629) in SV basal cells. In addition to regulation of ion homeostasis and pH, these analyses provide some insight into additional functions and processes in which SV cell types play a role.

#### Gene regulatory network extent appears related number of genes detected per cell or nuclei

scRNA-Seq and snRNA-Seq datasets are subject to a phenomenon called “dropout” or detection bias, where expression of a gene may be observed in one cell but is not detected in another cell of the same type (Kharchenko, Silberstein, & Scadden, 2014). This phenomenon results from a combination of the low amounts of mRNA in individual cells, inefficient mRNA capture, and the stochastic nature of mRNA expression. Because of this, scRNA-Seq and snRNA-Seq datasets exhibit a high degree of sparsity resulting in the detection of a small fraction of the transcriptome in a given cell. In comparing the effect of detection bias in our scRNA-Seq and snRNA-Seq datasets, we made several observations with respect to the ability to identify potential gene regulatory networks (GRNs) and identify clusters of cells.

In general, the number of genes detected per cell was greater than the genes detected per nuclei (Figure 6A). When examining genes detected amongst cell type-specific cells or nuclei (marginal, intermediate, or basal cells), this observation remained consistent (Figure 6A). In contrast, there was a greater number of genes that demonstrated significant differential expression among the three major cell types in the snRNA-Seq dataset (Figure 6B). Examining the gene expression amongst cell type-specific WGCNA modules (gene regulatory networks of co-expressed genes not linked by transcription factors) revealed a trend towards a greater number of genes detected in the scRNA-Seq dataset (Figure 6C). This observation became more definitive when examining gene expression amongst cell type-specific SCENIC regulons (gene regulatory networks of co-expressed genes linked by a common transcription factor) (Figure 6D). As a whole, these data suggest that gene regulatory network inference may be dependent on the number of genes detected per cell or nuclei and that nuclei-based approaches may allow for the detection of more differentially expressed genes in certain contexts.

**Figure 6.**
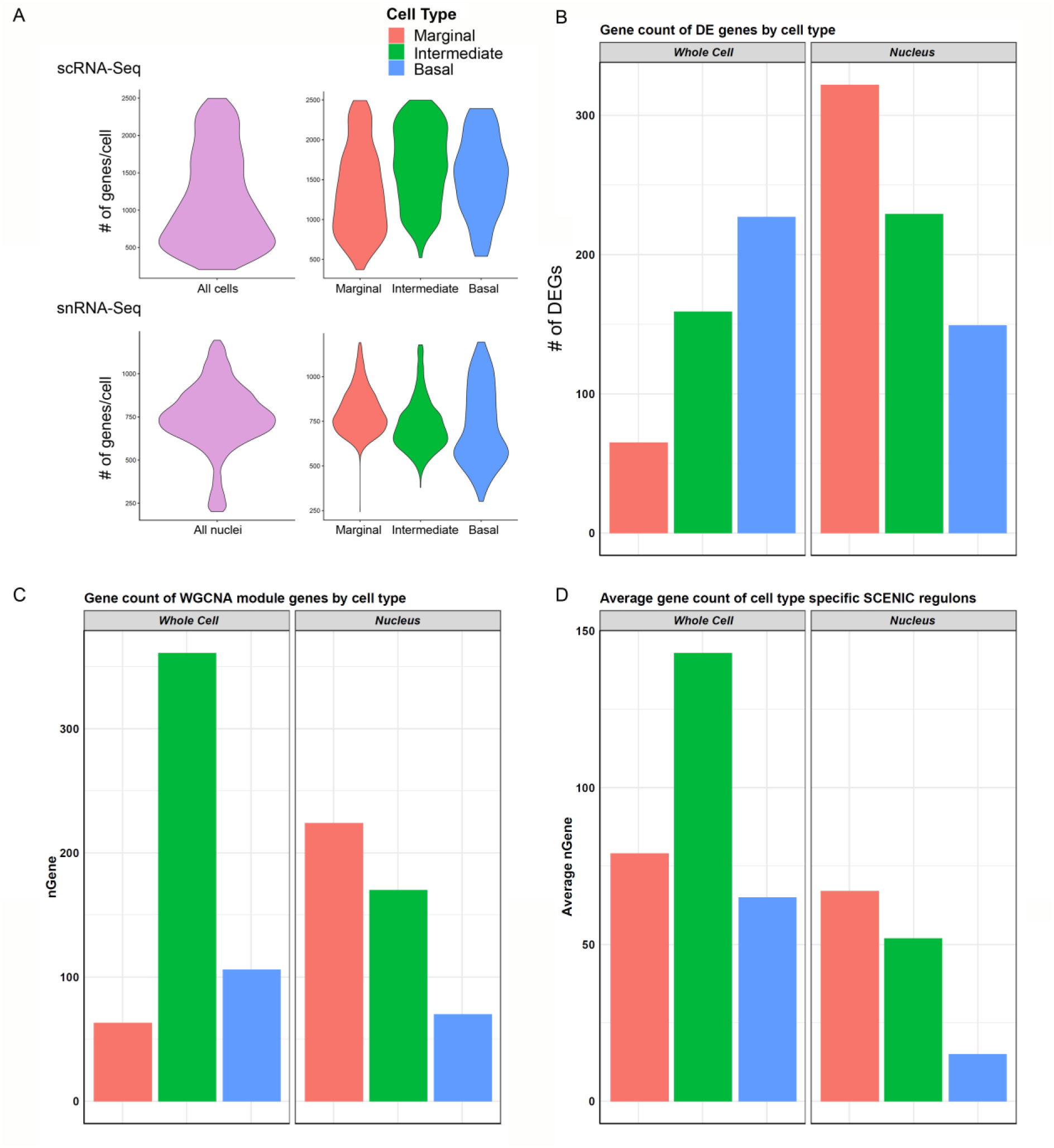
Cell type-specific gene expression comparison between scRNA-Seq and snRNA-Seq. **(A)**, Violin plots to left in upper (scRNA-Seq) and lower (snRNA-Seq) panels show the distribution of the number of genes per cell (upper panel) or nuclei (lower panel) along the vertical axis. In general, transcriptome profiles from cells exhibited a greater number of genes per cell than transcriptome profiles from nuclei. This trend continued as expected when looking at cell type-specific gene expression in marginal, intermediate, and basal cells in the panels to the right in both scRNA-Seq (upper panel) and snRNA-Seq (lower panel). **(B)** Cell type-specific expression of differentially expressed genes (DEGs) was lower in the scRNA-Seq data compared to the snRNA-Seq data in marginal, intermediate, and basal cells. nGene refers to number of differentially expression genes. **(C),** Gene counts for WGCNA cell type-specific modules did not demonstrate a consistent relationship between scRNA-Seq and snRNA-Seq datasets. **(D),** Gene counts for SCENIC cell type-specific regulons in general demonstrated a higher gene count per cell type in scRNA-Seq compared to snRNA-Seq. nGene (# of genes).

#### Validation of selected marginal cell- and intermediate cell-specific SCENIC regulons

In order to begin to identify critical gene regulatory networks, we identified cell type-specific SCENIC regulons identified in common between both the scRNA-Seq and snRNA-Seq datasets. We then examined these common regulons and utilized the regulon components identified in the single cell RNA-Seq dataset. Based on these parameters, two cell type-specific regulons were selected for further validation of the gene regulatory network inference approach. The estrogen-related receptor beta (*Esrrb*) regulon, a marginal cell-specific regulon, and the brain expressed myelocytomatosis oncogene (*Bmyc*), an intermediate cell-specific regulon, were selected for further analysis. Mutations in *Esrrb* result in autosomal recessive sensorineural hearing loss (DFNB35) (Collin et al, 2008) and missense mutations have recently been implicated in Meniere’s disease (Gallego-Martinez, Requena, Roman-Naranjo, & Lopez-Escamez, 2019). *Bmyc* has not previously been characterized as having a role in inner ear pathology or hearing loss. Selected regulons were screened for transcription factor targets with available smFISH probes, resulting in a small subset of genes from these regulons being selected.

Direct gene targets for *Esrrb* selected for validation with smFISH were *Abcg1*, *Heyl*, and *Atp13a5*. The gene activity plot shows the composite expression of the *Esrrb* regulon in SV marginal cells (Figure 7A). Binding motifs for *Esrrb* in each of the respective downstream targets (*Abcg1*, *Heyl*, *Atp13a5*) are shown (Figure 7B). Feature plots demonstrating RNA expression for *Abcg1* and *Heyl* have been previously shown in Figures 3A and 3C. Feature plots demonstrate RNA expression for *Esrrb* and *Atp13a5* in marginal cells (Figure 7C). smFISH demonstrates co-expression of *Esrrb* with each downstream target protein in SV marginal cells (Figure 7D).

**Figure 7.**
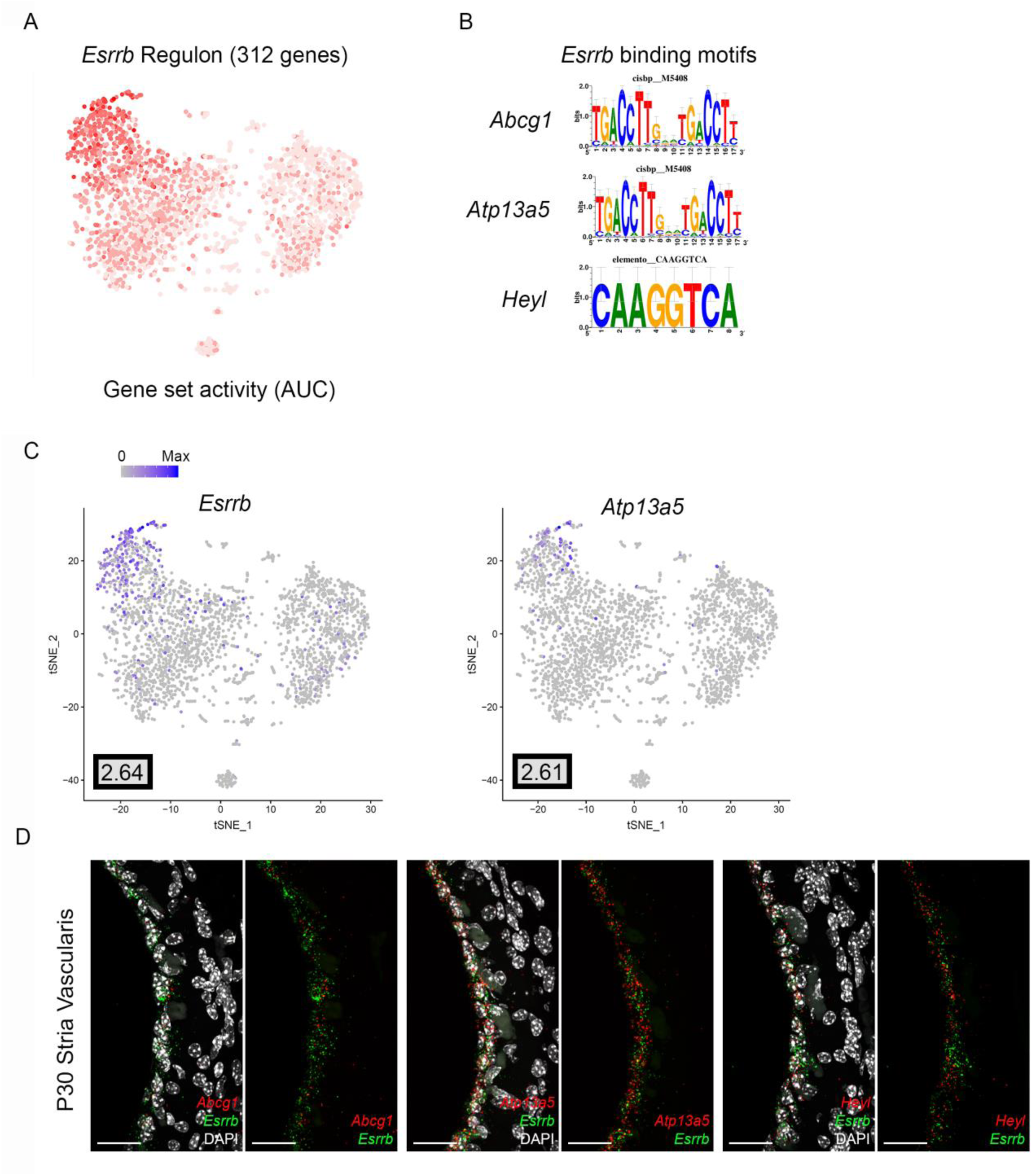
Validation of *Esrrb* regulon. **(A)**, Gene set activity plot displays *Esrrb* regulon expression in scRNA-Seq dataset. The more red a dot is, the greater the regulon activity in that given cell. **(B),** *Esrrb* binding motifs for *Esrrb* downstream targets (*Abcg1*, *Atp13a5*, *Heyl*). **(C),** Feature plots demonstrating expression of Esrrb and Atp13a5 in the marginal cell cluster in the scRNA-Seq dataset. Maximum expression is noted in the black outlined gray box for each gene in the lower left corner of each feature plot. Histogram is depicted from 0 to maximum (Max) in terms of normalized counts. Feature plots for *Abcg1* and *Heyl* have been shown previously in Figure 3. **(D),** smFISH demonstrates co-expression of *Abcg1*, *Atp13a5*, and *Heyl* transcripts in red with *Esrrb* transcripts in green in marginal cells on representative cross-sections of the P30 mouse SV. Panels to left depict *Esrrb* transcripts (red) and downstream target transcripts (green) with DAPI. Panels to the right depict *Esrrb* transcripts (red) and downstream target transcripts (green) without DAPI. DAPI (4′,6-diamidino-2-phenylindole). All scale bars 20 μm.

Direct gene targets for *Bmyc* selected for validation with smFISH were *Cd44*, *Met*, *Pax3*. The gene activity plot shows the composite expression of the *Bmyc* regulon in SV intermediate cells (Figure 8A). Binding motifs for *Bmyc* in each of the respective downstream targets (*Cd44*, *Met*, *Pax3*) are shown (Figure 8B). Feature plots demonstrating RNA expression for *Cd44* and *Met* have been previously shown in Figures 2B and 2D. Feature plots demonstrate RNA expression for *Bmyc* and *Pax3* in intermediate cells (Figure 8C). smFISH demonstrates *Bmyc* co-expression with each downstream target protein in SV intermediate cells (Figure 8D). Overall, the co-expression of transcription factors with their downstream targets demonstrates the strength of the SCENIC-based approach for gene regulatory network inference as well as clustering. The identification of regulons provides a starting point for targeted modulation of gene activity within a regulon to elucidate the role of the regulon in SV function.

**Figure 8.**
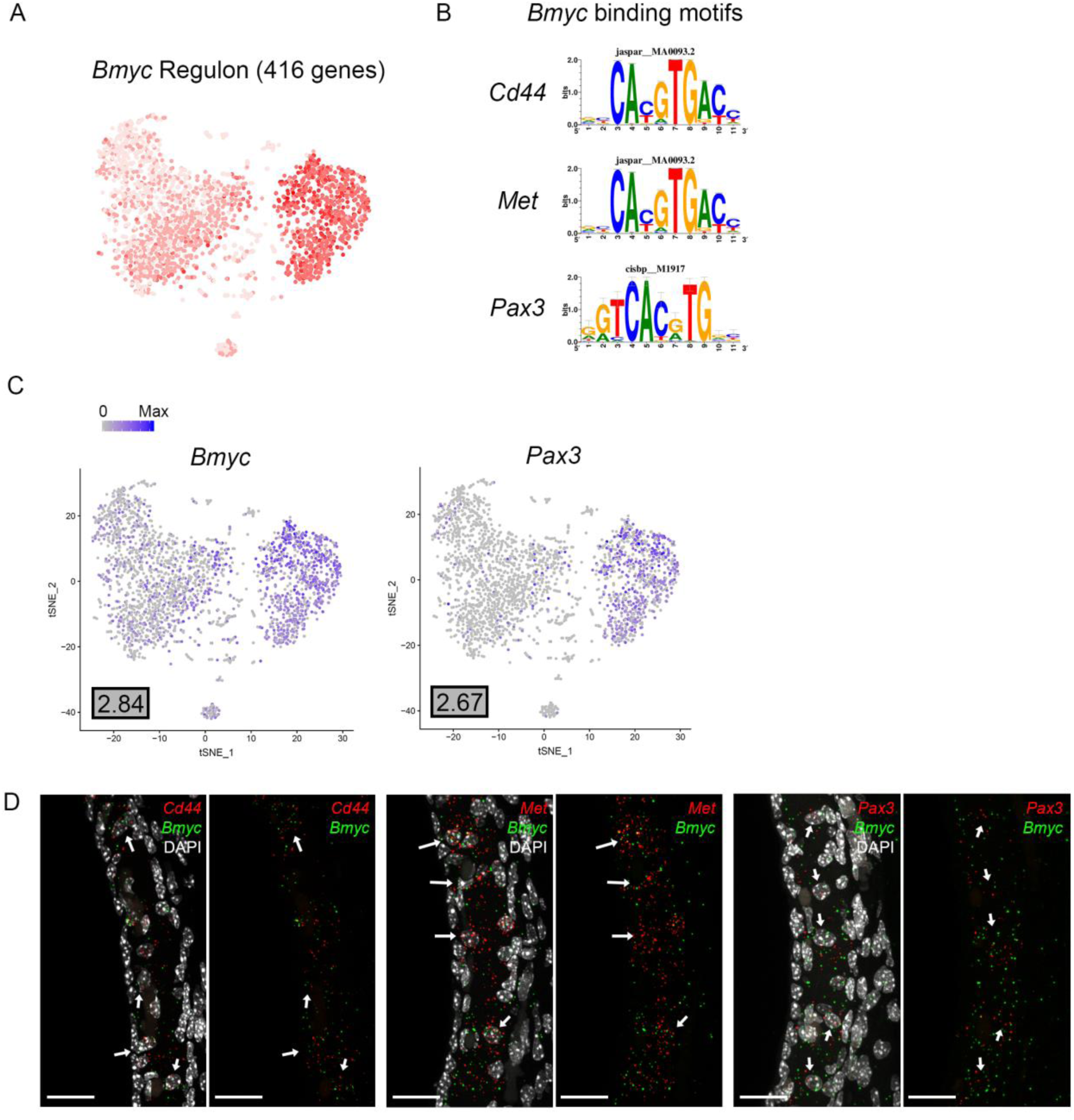
Validation of *Bmyc* regulon. **(A)**, Gene set activity plot displays *Bmyc* regulon expression in scRNA-Seq dataset. The more red a dot is, the greater the regulon activity in that given cell. **(B),** *Bmyc* binding motifs for *Bmyc* downstream targets (*Cd44*, *Met*, *Pax3*). **(C),** Feature plots demonstrating expression of *Bmyc* and *Pax3* in the intermediate cell cluster of the scRNA-Seq dataset. Maximum expression is noted in the black outlined gray box for each gene in the lower left corner of each feature plot. Histogram is depicted from 0 to maximum (Max) in terms of normalized counts. Feature plots for *Cd44* and *Met* have been shown previously in Figure 2. **(D),** smFISH demonstrates co-expression of *Cd44*, *Met*, and *Pax3* transcripts in red with *Bmyc* transcripts in green in intermediate cells on representative cross-sections of the P30 mouse SV. Panels to left depict *Bmyc* transcripts (red) and downstream target transcripts (green) with DAPI. Panels to the right depict *Bmyc* transcripts (red) and downstream target transcripts (green) without DAPI. DAPI (4′,6-diamidino-2-phenylindole). White arrows point to examples of intermediate cells that exhibit both Bmyc and downstream target transcript expression. All scale bars 20 μm.

#### Pharos analysis identifies druggable gene targets in cell type-specific regulons

With regards to modulation of gene activity within a regulon, Pharos, a database of druggable gene targets, provides an opportunity to identify FDA-approved drugs and small molecules that could potentially be utilized to modulate gene expression (https://pharos.nih.gov/). Analysis with Pharos identified FDA-approved drugs with known activities against cell type-specific regulon targets. The analysis was focused on Tclin targets, which are genes whose expression can be altered by a known mechanism of action of an FDA-approved drug. As an example of this approach, the previously described *Esrrb* and *Bmyc* regulons were utilized to construct a network diagram of their direct druggable targets. Direct druggable target genes of the *Esrrb* and *Bmyc* regulons are shown (Figure 9A & 9B, respectively). For the purposes of simplifying the display, activators included receptor agonists, enzyme activators, and ion channel openers. Inhibitors included receptor antagonists, receptor inverse agonists, enzyme inhibitors, and ion channel blockers. These drugs represent potential mechanisms by which strial function might be modulated.

**Figure 9.**
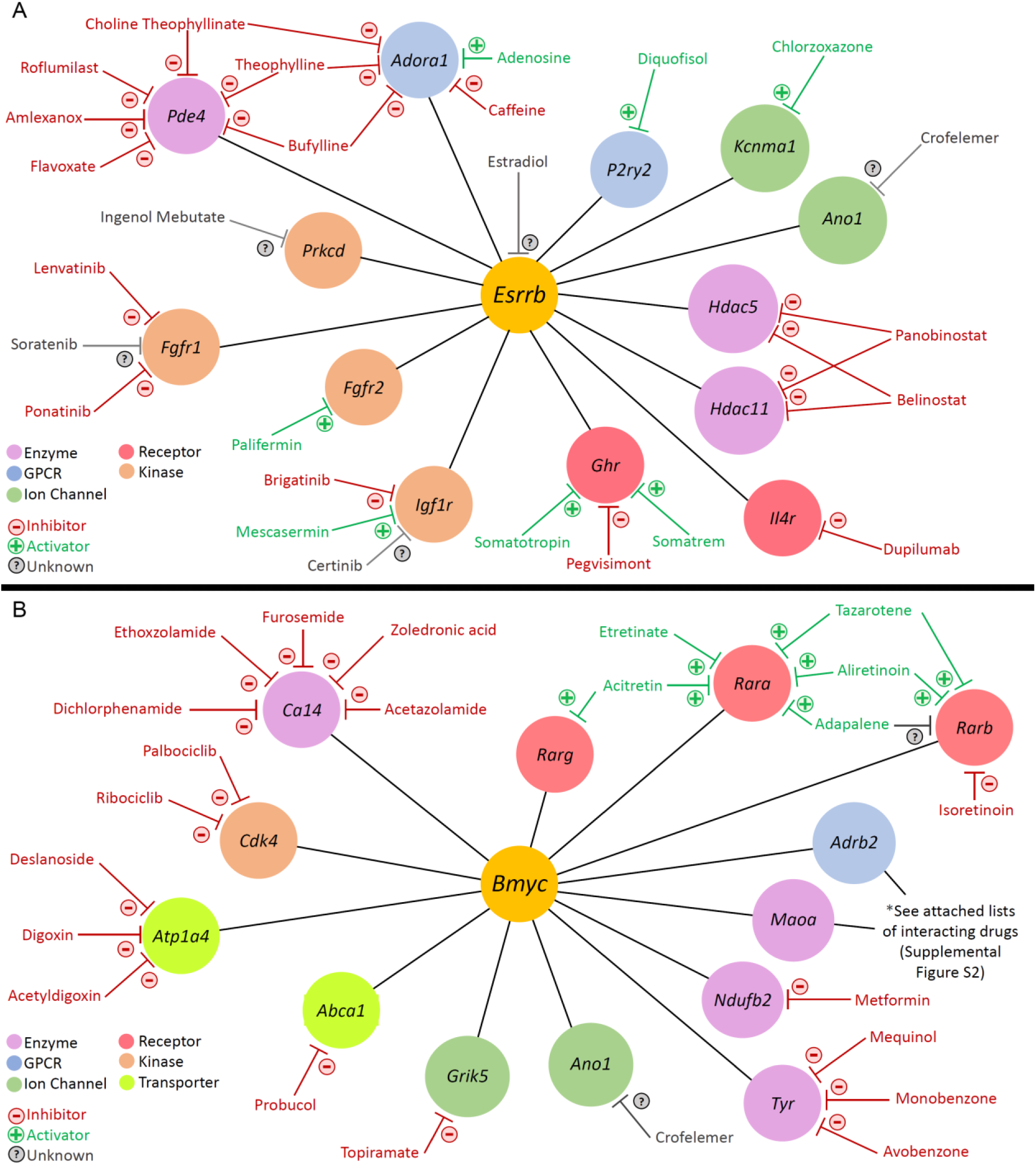
Analysis with Pharos reveals druggable gene targets in the *Esrrb* and *Bmyc* regulons. Pharmacologic drug classes are shown in lower left corner including enzyme, G-protein coupled receptor (GPCR), ion channel, receptor, and kinase. FDA-approved drugs are classified as inhibitors (red), activators (green), or unknown (grey). **(A),** Druggable gene targets in the *Esrrb* regulon. Analysis identified 13 druggable genes with 25 FDA-approved drugs. **(B),** Druggable gene targets in the *Bmyc* regulon. Analysis identified 13 druggable genes with 68 FDA-approved drugs. Drugs for *Adrb2* and *Maoa* genes are provided in Supplemental Figure S3.

For example, in the *Esrrb* regulon, studies suggest that estrogen (estradiol) may have a role in hearing protection (Milon et al., 2018; Wijayaratne & McDonnell, 2001; Williamson, Ding, Zhu, & Frisina, 2019). Milon and colleagues have suggested the possibility of utilizing estrogen signaling pathway effectors for hearing protection (Milon et al., 2018). In the *Bmyc* regulon, both metformin and acetazolamide have been associated with attenuation of hearing loss in certain settings (H. C. Chen, Chung, Lu, & Chien, 2019; Muri, Le, Zemp, Grandgirard, & Leib, 2019; Sepahdari, Vorasubin, Ishiyama, & Ishiyama, 2016; Wester, Ishiyama, Karnezis, & Ishiyama, 2018). Metformin, which inhibits *Ndufb2*, may reduce the incidence of sudden sensorineural hearing loss amongst diabetic patients (H. C. Chen et al., 2019; Muri et al., 2019). While the effects of acetazolamide in treating hearing loss associated with Meniere’s disease appear to be temporary (Crowson, Patki, & Tucci, 2016; Hoa, Friedman, Fisher, & Derebery, 2015), recent reports suggest some utility in sudden sensorineural hearing loss associated with PDE5 inhibitors (Wester et al., 2018).

### Deafness gene mapping suggests a role for SV cell types in human deafness

Finally, we screened the cellular transcriptomes from our scRNA-Seq and snRNA-Seq datasets for genes associated with human hearing loss. The database of deafness genes was constructed from resources including the Hereditary Hearing Loss Homepage as previously described previously (Azaiez et al., 2018; Shearer, Hildebrand, & Smith, 1993). Heatmaps showing gene expression for known deafness genes in the P30 stria vascularis are shown in Figure 10. Expression data corresponding to P30 SV scRNA-Seq is shown in the left heatmap (Figure 10A) and expression data corresponding to P30 SV snRNA-Seq is show in the right heatmap (Figure 10B). Known deafness genes are displayed along the vertical axis and SV cell types are displayed horizontally with marginal cells in pink, intermediate cells in green, basal cells in blue, and spindle cells in yellow.

**Figure 10.**
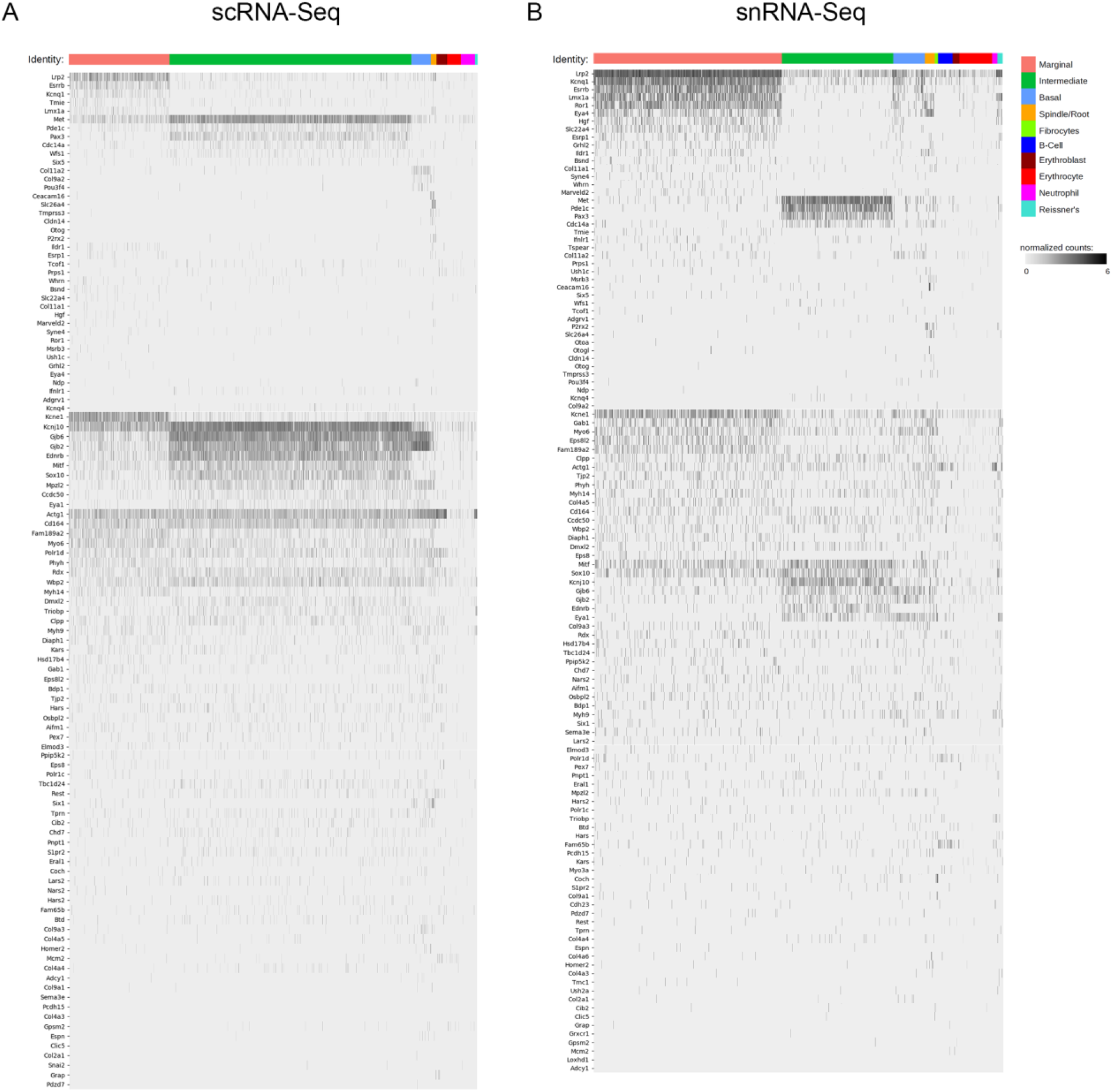
Deafness gene mapping of adult stria vascularis single cell and single nucleus transcriptomes associates SV cell types with deafness genes. Cell types are arrayed along the horizontal axis and grouped according to cell types. Deafness genes are arrayed along the vertical axis. Deafness genes were grouped according to how specific their expression seemed to be in a given cell type. Genes with less cell type-specific expression were placed lower in the list of genes. **(A),** Expression level in normalized counts for each deafness gene (each row) across all cells in the adult SV scRNA-Seq dataset. **(B),** Expression level in normalized counts for each deafness gene (each row) across all cells in the adult SV snRNA-Seq dataset.

We identify a subset of deafness genes that are expressed in adult SV cell types. Deafness genes expressed in marginal cells include *Lrp2*, *Esrrb*, *Hgf*, *Kcnq1*, *Kcne1*, *Ror1*, and *Eya4* (Collin et al., 2008; Faridi et al., 2019; Khalifa et al., 2015; König et al., 2008; Schultz et al., 2009; Wangemann, 2002; Wayne, 2001; Zhang, Knosp, Maconochie, Friedman, & Smith, 2004). Deafness genes expressed in intermediate cells include *Met*, *Pde1c*, and *Pax3* (Bondurand, 2000; Mujtaba et al., 2015; Tassabehji et al., 1993; Wang et al., 2018). *Col11a2* appears to be specifically expressed in SV basal cells (McGuirt et al., 1999). In addition to *Slc26a4*, deafness genes expressed in spindle cells include *Ceacam16*, *Cldn14*, and *P2rx2* (Ben-Yosef et al., 2003; Booth, Kahrizi, Najmabadi, Azaiez, & Smith, 2018; Faletra et al., 2014; Naz et al., 2017; Wilcox et al., 2001; Zheng et al., 2011; Zhu, Beudez, Yu, Grutter, & Zhao, 2017). More importantly, these data demonstrate the possibility that some of these deafness genes affect multiple SV cell types, as in the case of *Mitf* and *Sox10,* which are expressed in both intermediate and marginal cells (Bondurand, 2000). Both genes are expressed by neural crest cells, suggesting that multiple cell types in the stria vascularis may arise from neural crest cells. The contribution of neurocristopathies to hearing loss is a developing area of research (Hao et al., 2014; Locher et al., 2015; Ritter & Martin, 2019; Shibata et al., 2016). Another example of hearing loss where multiple cell types may be affected includes Connexin 26 and 30 encoded by *Gjb2* and *Gjb6*, respectively, which appear to be expressed in intermediate and basal cells and, to a lesser extent, marginal cells (Figure 10) (W. Liu, Boström, Kinnefors, & Rask-Andersen, 2009; Mei et al., 2017). These data reveal expression of a subset of human deafness genes in SV cell types and define potential opportunities for further investigation into the function of these genes in the SV.

## Discussion

Dual methodologies were used to develop an atlas of adult SV cell type transcriptional profiles, the most comprehensive to date. Specifically, the use of dual RNA-Seq methodologies (scRNA-Seq and snRNA-Seq) revealed similar adult SV cell type-specific transcriptional profiles. Furthermore, we demonstrate the utility of applying dual gene regulatory network inference methodologies to both corroborate cell type heterogeneity as well as define potentially critical gene regulatory networks. We provide examples of how this resource can be harnessed to identify and confirm cell type-specific gene regulatory networks, identify druggable gene targets, and implicate SV cell types in syndromic and nonsyndromic hereditary deafness. We now define the implications of these data for: (1) using and developing mouse models to define the contribution of each SV cell type to SV function and hearing at the level of gene expression, (2) understanding ion homeostatic mechanisms broadly in the inner ear, (3) contributing to efforts to develop targeted SV gene therapy through the use of cell type-specific promoters, (4) identifying druggable gene targets and the potential for repurposing existing pharmacotherapies to treat inner ear disease, and (5) use of these data to identify cell type-specific contributions to disease.

### Implications for mouse model development to enhance our understanding of cell type-specific mechanisms related to the loss of EP and hearing loss

In characterizing adult SV cellular heterogeneity using transcriptional profiling, these data provide opportunities to study adult SV cell type-specific mechanisms that contribute to EP generation and whose dysfunction results in hearing loss. Novel cell type-specific gene expression allows for the identification of existing inducible cell type-specific Cre-recombinase mice as well as the development of novel inducible mouse models to study of the effects of gene deletion on adult SV cell types that can be separated from developmental effects. Genetic mutations that affect the SV result in hearing loss that is characterized by an initial loss of EP followed by a delayed loss of hair cells (Huebner et al., 2019; H. Liu et al., 2016). These existing mouse models are complicated by the developmental effects of genes critical to EP generation (Huebner et al., 2019; H. Liu et al., 2016). Conversely, mouse models of SV age-related hearing loss are complicated by incompletely defined mechanisms and time windows which make for challenging experimental conditions (Ohlemiller, 2009). Thus, the exact timing and mechanisms associated with a permanent loss of hair cells that follow the loss of EP remain incompletely defined. Use of inducible mouse models may enable insight into the timing of these mechanisms as well as insight into possible mechanisms. Understanding the critical time window for intervention as well as potential targeted interventions that might slow, halt or reverse the effects of EP loss will contribute to development of therapeutic approaches to hearing loss due to SV dysfunction.

### Implications for understanding ionic homeostasis in the inner ear

SV cell type-specific gene regulatory networks include genes critical to EP generation and ion homeostasis and as a result provide a window into inner ear ion homeostatic mechanisms. Insight into these ion homeostatic mechanisms may be important to understanding their functional contribution to hearing and hearing loss (Nickel & Forge, 2008; Zdebik, Wangemann, & Jentsch, 2009). These gene regulatory networks within the unperturbed wild type adult mouse SV establish a basis for identifying and interpreting changes in these networks in disease settings including, but not limited to, Meniere’s disease (Collin et al., 2008; Ishiyama, Tokita, Lopez, Tang, & Ishiyama, 2007; Kariya et al., 2009), autoimmune inner ear disease (Calzada, Balaker, Ishiyama, Lopez, & Ishiyama, 2012), cisplatin-induced hearing loss (Breglio et al., 2017; Rybak, Mukherjea, & Ramkumar, 2019), and enlarged vestibular aqueduct syndrome (T. Ito, Nishio, Wangemann, & Griffith, 2015; Miyagawa et al., 2014).

While the consequences of dysfunctional ion homeostasis to human hearing loss are incompletely understood, ion homeostasis appears to be critical, as evident by the number of deafness genes encoding ion channels and transporters, many of which are expressed in the SV (Mittal et al., 2017). While this study demonstrates expression of these ion channels in SV cells in the cochlea, there are quite a few examples of marginal cell-specific genes that are also expressed in the vestibular portion of the inner ear (Bartolami et al., 2011; Jichao Chen & Nathans, 2007; Wangemann, 1995). The consequences of dysfunctional ion homeostasis on vestibular function remain incompletely characterized (Jones & Jones, 2014). Insights gained in understanding the mechanisms regulating ion homeostasis in the SV may apply to other regions of the inner ear. The co-occurrence of hearing loss and renal disease resulting from mutations in channel and transporter genes suggests that understanding ion homeostasis in the inner ear may have broader implications to dysfunctional ion homeostasis and human disease (Lang, Vallon, Knipper, & Wangemann, 2007).

### Implications for targeting gene therapy to SV cell types

Cell type-specific genes could be utilized to create viral vectors with cell type-specific promoters for improved therapeutic targeting to the SV. This idea is not without precedent as cell type-specific promoters have been utilized in the brain, retina and inner ear to target specific groups of cells (Ingusci, Verlengia, Soukupova, Zucchini, & Simonato, 2019; Kim et al., 2013; McDougald et al., 2019; Praetorius et al., 2010). Furthermore, some recent adoptive immunotherapy trials using a cell type-specific promoter for melanoma already suggest the possibility for this in relation to the stria vascularis (Duinkerken et al., 2019; Johnson et al., 2009; Seaman et al., 2012). Specifically, these studies utilized adoptive immunotherapy using T cell receptors (TCRs) targeting MART-1, a known melanoma-associated antigen, which resulted in sensorineural hearing loss that in some cases was ameliorated by local steroid administration (Johnson et al., 2009). SV melanocytes arise from neural crest cells that migrate into the SV during development (Shibata et al., 2016). SV melanocytes represent the future intermediate cells (Shibata et al., 2016; Steel & Barkway, 1989). These studies suggest the possibility that vectors utilizing the gene that encodes MART-1, Melan-A (*Mlana*), may be an effective promoter for targeting gene therapy to SV intermediate cells.

Alternatively, gene therapy for nonsyndromic sensorineural hearing loss may entail targeting multiple cell types. For example, we demonstrate the expression of *Gjb2* and *Gjb6*, which encodes connexin 26 and connexin 30 proteins, respectively, in SV intermediate, basal and to a lesser extent, marginal cells (Fig. 10). Mutations in *Gjb2* and *Gjb6* followed by digenic mutations in these genes are among the most common causes of nonsyndromic autosomal recessive sensorineural hearing loss (SNHL) in many populations across the world (Mei et al., 2017; Nickel & Forge, 2008). Mei and colleagues demonstrate that the loss of *Gjb2* and *Gjb6* in the stria vascularis and lateral wall result in a loss of EP and hearing loss while the loss of *Gjb2* and *Gjb6* in cochlear supporting cells does not (Mei et al., 2017). These data identify the loss of these genes in the SV as principal drivers of hearing loss in digenic *Gjb2* and *Gjb6* mutations. Use of SV promoters that may be capable of targeting multiple SV cell types including *Mitf* or *Sox10* could be utilized in future therapeutic attempts (Fig. 10) (Hao et al., 2014; Walters & Zuo, 2015).

### Implications for pharmacologic targeting of SV cell types

Our analyses utilizing parallel gene regulatory network inference methods (WGCNA and SCENIC) combined with a search for druggable gene targets utilizing Pharos (https://pharos.nih.gov/) suggest a possible translational approach to utilizing scRNA-Seq and snRNA-Seq datasets (Aibar et al., 2017; Langfelder & Horvath, 2008; Nguyen et al., 2017). Through this approach, we identify genes for which FDA-approved drugs have a known effect. This agnostic in silico approach can potentially be utilized to identify FDA-approved medications that could be repurposed or repositioned to treat diseases in the inner ear as has been done in other areas of human disease including neurodegenerative diseases, cancer, and autoimmune disease (Ferrero & Agarwal, 2018; Paranjpe, Taubes, & Sirota, 2019). This may be particularly useful in hearing and balance disorders that are characterized by onset of symptoms in adulthood (i.e. autoimmune inner ear disease, Meniere’s disease). We acknowledge that there are many methods to identify drugs for repurposing or repositioning, including in silico methods, which have been reviewed elsewhere (Ferrero & Agarwal, 2018; Li et al., 2016; Vanhaelen et al., 2017). However, our analyses highlight the potential use of the data for drug repurposing approaches.

### Implications for identifying cell type-specific contributions to disease

Finally, as we have alluded to previously, scRNA-Seq and snRNA-Seq approaches could potentially be utilized to associate changes in gene expression in human disease with cell types in a given tissue (Skene & Grant, 2016). Skene and colleagues associated cell types with human disease by comparing human disease expression datasets for Alzheimer’s disease, autism, schizophrenia, and multiple sclerosis to single cell transcriptional profiles from the mouse cortex and hippocampus. In a similar fashion, single cell and single nucleus transcriptome profiles, like the resource we have developed for the adult stria vascularis, might be utilized to associate cell types to inner ear diseases. For example, in Meniere’s disease, while no single gene has been conclusively proven to cause disease, the association of gene mutations with cell type-specific expression might provide some clues to the involved cell type or types (Chiarella, Petrolo, & Cassandro, 2015; Lopez-Escamez, Batuecas-Caletrio, & Bisdorff, 2018). Mutations in both *Kcne1* and *Esrrb* have been identified in patients with Meniere’s disease (Dai, Wang, & Zheng, 2019; Gallego-Martinez et al., 2019; Lopes, Sartorato, Da Silva-Costa, De Macedo Adamov, & Ganança, 2016). Our data demonstrates expression of these genes in SV marginal cells (Fig. 10). It is possible that marginal cells might play a role in the pathophysiology of this disease.

While this is the most comprehensive cell atlas of the adult SV to date, some limitations or caveats are worth mentioning. We did not definitively identify clusters of pericytes or endothelial cells in our dataset and chose to focus our analysis on the four main groups of cells (marginal, intermediate, basal, and spindle cells). Given the lower number of spindle and root cells, we did not definitively distinguish spindle and root cells from each other. Finally, the scRNA-Seq dataset consisted of 3 separate captures. These runs were compared across clusters and demonstrated equal distribution across all cell clusters identified. For this reason, we combined these datasets and they are presented as one dataset.

We believe that the utility of these datasets as resources is expansive. These datasets establish a baseline for comparison to effects on the SV related to treatment and provide a resource for identifying cell types associated with human inner ear disease. We show examples of applications of these data, including characterization of homeostatic gene regulatory networks, druggable gene target analysis with Pharos to identify potential repurposing of FDA-approved medications, and provide the most focused screen of deafness gene expression in the SV. These data will serve as a baseline for identifying key regulatory mechanisms related to genetic and acquired hearing loss, as well as, for responses to a variety of pharmacologic treatments.

## Materials and Methods

**Table 1.**
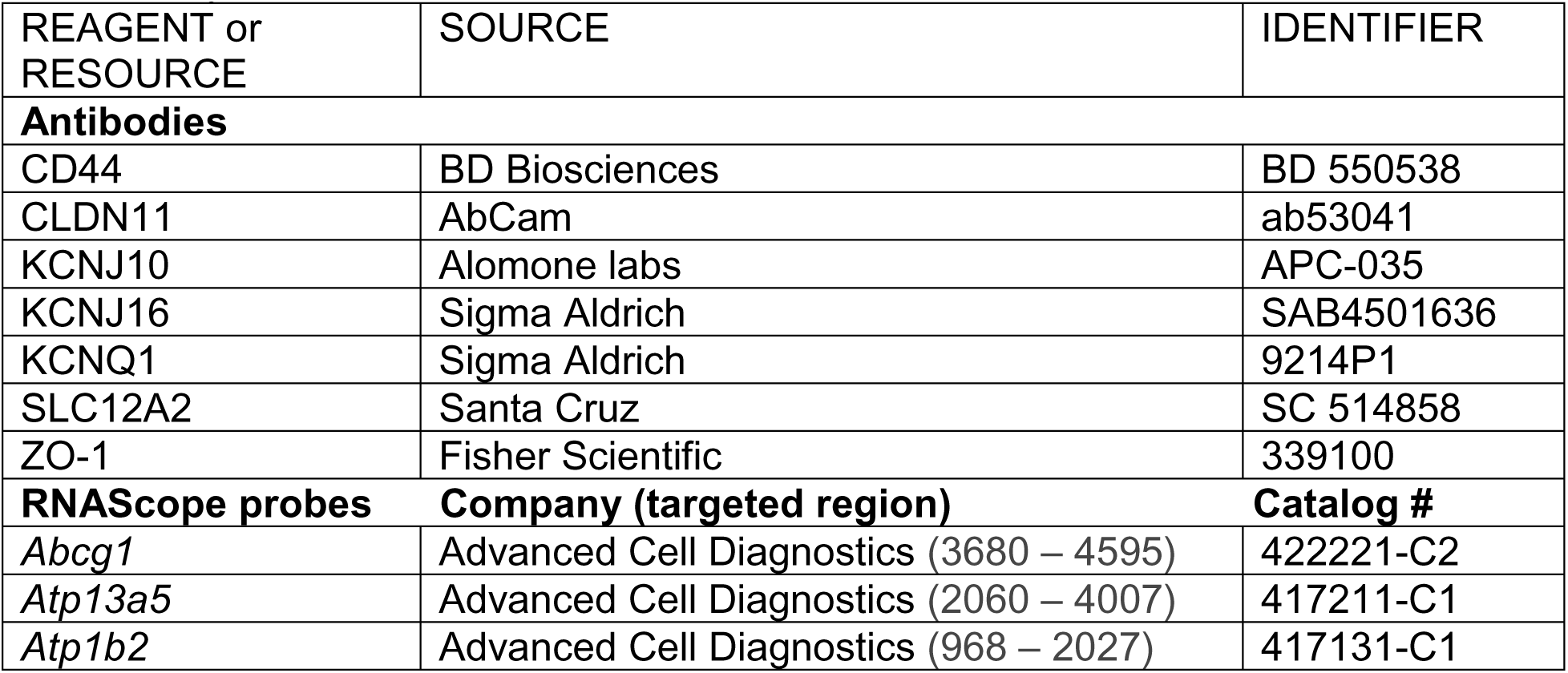
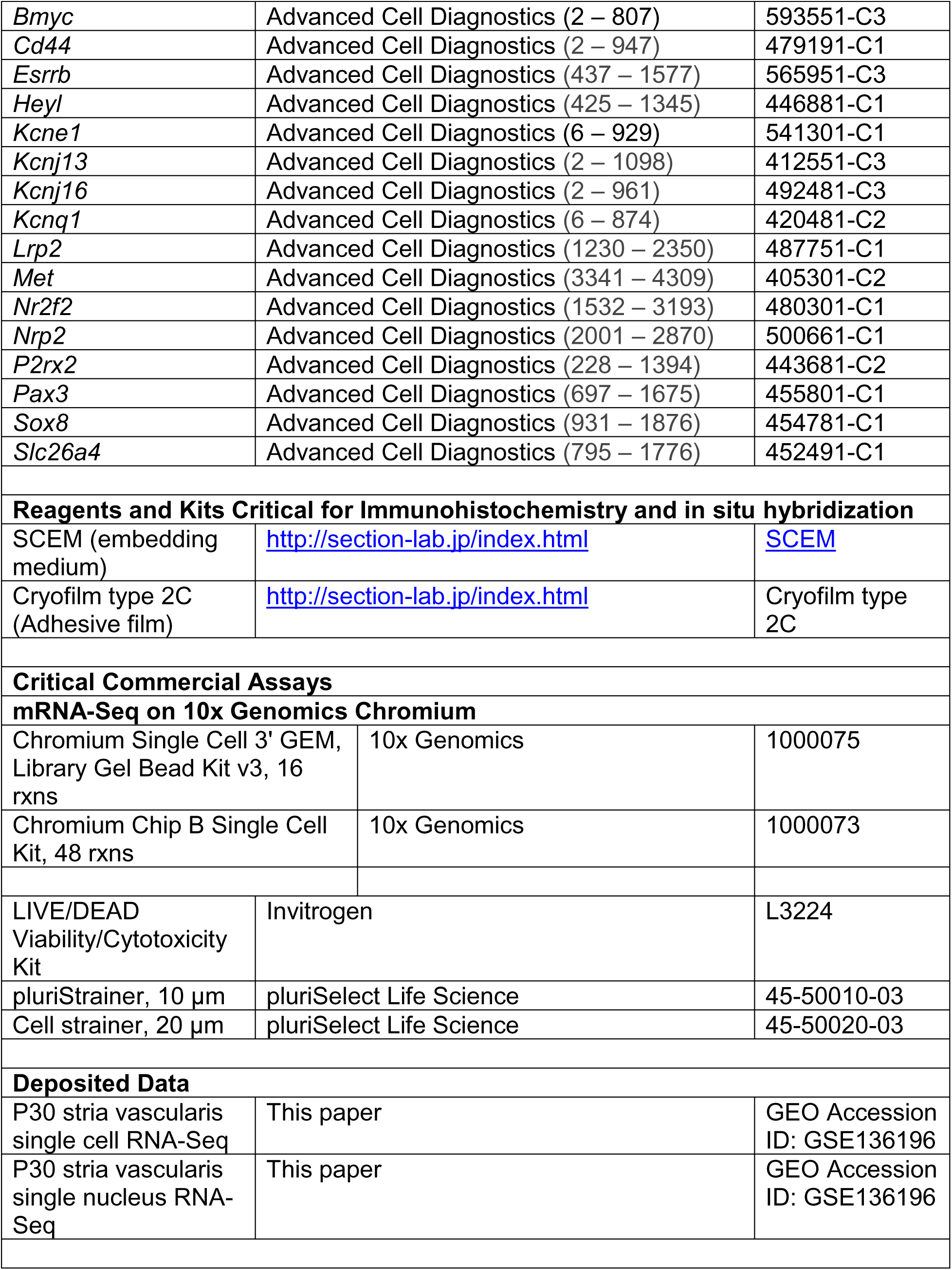
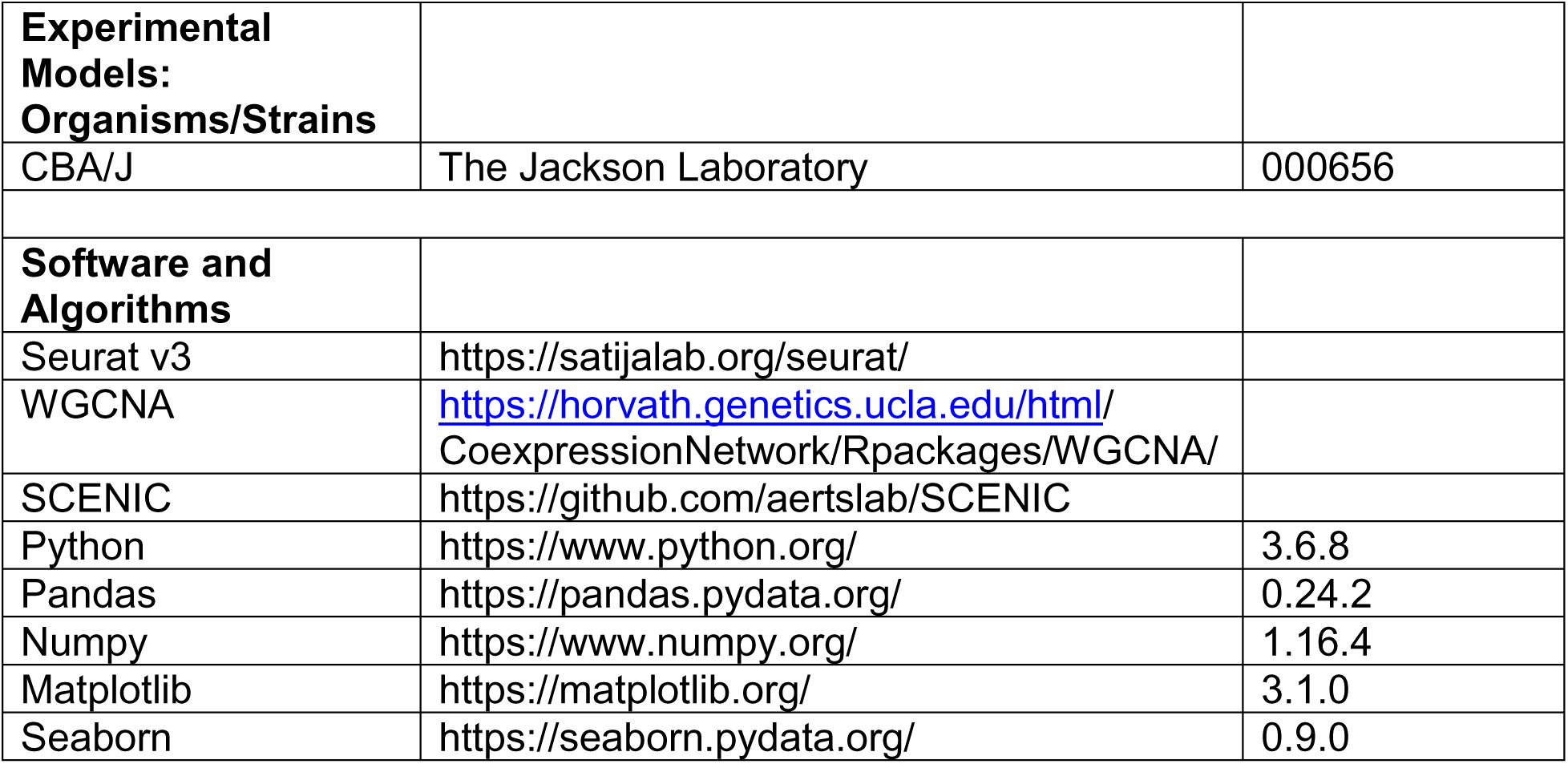
Key Resources

### Animals

Inbred CBA/J males and females were purchased from JAX (Stock No. 000656). Breeding pairs were set up to obtain P30 mice for single cell and single nucleus RNA seq experiments, immunohistochemistry and single molecule RNA FISH.

### Cell/nucleus Isolation

#### Single cell suspension

Inner ears from a total of ∼25 P30 mice were removed and stria vascularis from the cochleae were collected into 200ul DMEM F-12 media. The tissue was lysed in 0.5mg/ml trypsin at 37c for 7 minutes. The media was carefully removed and replaced with 5% FBS to stop the lysis. The tissue was triturated and filtered through a 20um filter (pluriSelect Life Science, El Cajon, CA). The filtered cells were let to sit on ice for 35 minutes. 150ul of the supernatant was removed and cell pellet was suspended in the remaining 50ul. Cells were counted on a Luna automated cell counter (Logos Biosystems, Annandale, VA) and a cell density of 1×10^6^ cells/ml was used to load onto the 10X genomics chip.

#### Single nucleus suspension

Published single nucleus suspension protocol from 10x genomics was used to isolate the nuclei. Briefly, stria vascularis from ∼10 P30 animals were isolated and collected in 3ml DMEM F-12 media. Following collection, the media was replaced with 3ml chilled lysis buffer (10mM Tris-HCl, 10mM Nacl, 3mM MgCl2,0.005% Nonidet P40 in Nuclease free water) and the tissue were lysed at 4c for 25 minutes. The lysis buffer was then replaced with 1.5ml DMEM F-12 media. The tissues were triturated and filtered through a 20um filter (pluriSelect Life Science, El Cajon, CA). The filtrate was centrifuged at 500rcf for 5 minutes at 4c. The supernatant was removed, and the cell pellet was resuspended in 1ml nuclei wash and resuspension buffer (1xPBS with 1% BSA and 0.2U/ul RNase Inhibitor). The cells were filtered through a 10um filter (pluriSelect Life Science, El Cajon, CA) and centrifuged at 500rcf for 5 minutes at 4c. The supernatant was removed, and pellet resuspended in 50ul of nuclei wash and resuspension buffer. Nuclei were counted in a Luna cell counter (Logos Biosystems, Annandale, VA) and a nuclear density of 1×10^6^ cells/ml was used to load onto the 10X genomics chip.

#### 10x Chromium genomics platform

Single cell or nuclei captures were performed following manufacturer’s recommendations on a 10x Genomics Controller device (Pleasanton, CA). The targeted number of captured cells or nuclei ranged from 6,000 to 7,000 per run. Library preparation was performed according the instructions in the 10x Genomics Chromium Single Cell 3’ Chip Kit V2. Libraries were sequenced on a Nextseq 500 instrument (Illumina, San Diego, CA) and reads were subsequently processed using 10x Genomics CellRanger analytical pipeline using default settings and 10x Genomics downloadable mm10 genome. Dataset aggregation was performed using the cellranger aggr function normalizing for total number of confidently mapped reads across libraries.

### PCA and *t*-SNE Analysis

#### Selection of genes for clustering analysis

Identification of the highly variable genes was performed in Seurat utilizing the MeanVarPlot function using the default settings with the aim to identify the top ∼ 2000 variable genes (Satija, Farrell, Gennert, Schier, & Regev, 2015). Briefly, to control for the relationship between variability and average expression, average expression and dispersion is calculated for each gene, placing the genes into bins, and then a z-score for dispersion within the bins is calculated. These genes are utilized in the downstream analysis for clustering.

#### Clustering of single cells

Clustering analysis of single-cell data was performed with Seurat using a graph-based clustering approach (Satija et al., 2015). Briefly, the Jackstraw function using the default settings was used to calculate significant principal components (p < 0.0001) and these principal components were utilized to calculate k-nearest neighbors (KNN) graph based on the euclidean distance in PCA space. The edge weights are refined between any two cells based on the shared overlap in their local neighborhoods (Jaccard distance). Cells are then clustered according to a smart local moving algorithm (SLM), which iteratively clusters cell groupings together with the goal to optimize the standard modularity function (Blondel, Guillaume, Lambiotte, & Lefebvre, 2008; Waltman & van Eck, 2013)(http://satijalab.org/seurat/pbmc-tutorial.html). Resolution in the FindClusters function was set to 0.8. High modularity networks have dense connections between the nodes within a given module but sparse connections between nodes in different modules. Clusters were then visualized using a t-distributed stochastic neighbor embedding (t-SNE) plot.

#### Doublet detection and elimination

To detect and eliminate doublets that may affect clustering analysis we used an R package called DoubletDecon described previously (DePasquale et al., 2018) that iteratively detects doublets in single-cell RNASeq data. An expression matrix of read counts, the top 50 marker genes for each cluster, and a list of cells with cluster identities are used to create “medoids” for each cluster which are averages of cell-type specific gene expression. It then compares the similarity of expression for each cluster medoid to every other medoid resulting in a binary correlation matrix. If two cluster medoids meet or exceed the similarity threshold they receive a “1” in the binary matrix, referred to as “blacklist clusters”, and if they do not they receive a “0” in the binary matrix. Synthetic doublets are generated using cells from each pairwise comparison of dissimilar cluster medoids. Synthetic doublets, cells, and blacklist cluster gene expression profiles are deconvoluted and each cell is iteratively compared to both synthetic doublet profiles and blacklist clusters. If a cell is more similar in expression to synthetic doublets it is classified as a putative doublet. Putative doublets are re-clustered, and Welch’s t-test is used to determine uniquely expressed genes. If any cells express one unique gene when compared to the blacklisted clusters they are rescued and classified as a singlet. The singlet list was used to subset our single-cell data, and the Seurat pipeline was re-run on the new doublet eliminated data.

#### Differential gene expression analysis

Differential expression analysis was performed in Seurat utilizing the FindAllMarkers function with the default settings except that the “min.pct” and “thresh.use” parameters were utilized to identify broadly expressed (min.pct = 0.8, thresh.use = 0.01) and subpopulation-specific (min.pct = 0.5, thresh.use = 0.25) expression profiles. The parameter “min.pct” sets a minimum fraction of cells that the gene must be detected in all clusters. The parameter “thresh.use” limits testing to genes which show, on average, at least X-fold difference (log-scale) between groups of cells. The default test for differential gene expression is “bimod”, a likelihood-ratio test (McDavid et al., 2013). Differentially expressed genes were then displayed on violin plots based on unbiased clustering described above.

#### Heatmap or grid violin plot construction of selected data

Heatmaps or grid violin plots were constructed using custom python scripts and utilized to display gene expression across SV cell types identified in both scRNA-Seq and snRNA-Seq datasets. Briefly, construction of heatmaps or grid violin plots was performed in the following fashion: (1) Raw counts data was processed within Seurat as previously described. (2) Normalized counts were scaled into range [0,1] by using min-max scaling method on each gene. (3) Heatmaps or violin plots were constructed by python/seaborn using the scaled data counts. Detailed code can be found in python scripts in supplementary material.

### Downstream Computational Analysis

#### Gene regulatory network inference

Two independent methods of gene regulatory network inference, Weighted gene co-expression network analysis (WGCNA) (Langfelder & Horvath, 2008) and single cell regulatory network inference and clustering (SCENIC) (Aibar et al., 2017) were utilized. Briefly, WGCNA constructs a gene co-expression matrix, uses hierarchical clustering in combination with the Pearson correlation coefficient to cluster genes into groups of closely co-expressed genes termed modules, and then uses singular value decomposition (SVD) to determine similarity between gene modules. Hierarchical clustering of modules is displayed as topological overlap matrix (TOM) plots and similarity between gene modules are displayed as adjacency plots. Briefly, SCENIC identifies potential gene regulatory networks by performing the following steps: (1) ***modules*** consisting of transcription factors and candidate target genes are determined on the basis of co-expression utilizing GENIE3, (2) ***regulons*** are constructing by filtering modules for candidate genes that are enriched for their transcription factor binding motifs utilizing RcisTarget, (3) the activity of each regulon within each cell is determined, (4) the ***regulon activity matrix*** is constructed utilizing these regulon activity scores and can be used to cluster cells on the basis of shared regulatory networks. In this way, SCENIC may identify cell types and cell states on the basis of shared activity of a regulatory subnetwork.

#### Deafness gene screen of stria vascularis transcriptomes from scRNA-Seq and sn-RNA-Seq datasets

In order to screen our datasets for known deafness genes, we constructed a database of known human and mouse deafness genes from the following sources: (1) Hereditary Hearing Loss page (https://hereditaryhearingloss.org/)(Van Camp, Guy; Smith, n.d.) and (2) Hereditary hearing loss and deafness overview (Azaiez et al., 2018; Shearer et al., 1993).

#### Gene ontology, gene-set enrichment analysis and pathway-enrichment analysis of predicted proteins

Gene ontology analysis and gene enrichment analysis were performed using Enrichr (http://amp.pharm.mssm.edu/Enrichr/) as previously described (E. Y. Chen et al., 2013; Kuleshov et al., 2016; Pazhouhandeh et al., 2017). Enrichr is an integrated web-based application that includes updated gene-set libraries, alternative approaches to ranking enriched terms, and a variety of interactive visualization approaches to display the enrichment results. Enrichr employs three approaches to compute enrichment as previously described (Jagannathan et al., 2017). The combined score approach where enrichment was calculated from the combination of the p-value computed using the Fisher exact test and the z-score was utilized. In order to visualize molecular interaction networks, the list of putative proteins inferred from genes expressed by adult cochlear supporting cells was introduced to STRING 10.0 (http://string-db.org) and the nodes (proteins) and edges (protein-protein interactions) were extracted (Szklarczyk et al., 2015). Proteins were linked in STRING based on the default medium (0.400) minimum interaction score and on the following seven criteria: textmining, experiments, databases, co-expression, neighborhood, gene-fusion, and co-occurrence. Interaction evidence from all utilized criteria are benchmarked and calibrated against previous knowledge, using the high-level functional groupings provided by the manually curated Kyoto Encyclopedia of Genes and Genomes (KEGG) pathway maps (Szklarczyk et al., 2015). The summation of this interaction evidence is utilized to construct a minimum interaction score.

#### Identification of potentially druggable gene targets

To identify druggable targets within our scRNA-Seq data of the stria vascularis, genes from P30 cell-type specific SCENIC regulons were input into Pharos (https://pharos.nih.gov) (Nguyen et al., 2017) using their “batch search” function. Pharos is a database created by the “Illuminating the Druggable Genome” program to give users access to protein targets and the availability of drugs or small molecules for each. Pharos categorizes each protein with a “target developmental level” according to how much is known on its “druggability”. Targets that are well studied are deemed “Tclin” if they can be targeted with FDA approved drugs that have a known mechanism of action on the target, “Tchem” if there are known small-molecule ligands that bind the target, or “Tbio” if the target has a known gene ontology or phenotype but no available drugs or small molecules. Targets that are currently unstudied are labeled “Tdark”. To focus on the most clinically relevant targets, we filtered for only Tclin and Tchem developmental levels in our search. Tclin and Tchem targets from each cell-type specific regulon were plotted using the FeaturePlot function in Seurat to identify the most specific targets within regulons of the stria vascularis.

#### Fluorescent *in situ* hybridization (smFISH) using RNAscope probes

*In situ* hybridizations were performed using the following RNAscope probes (Table 1, Supplemental Table S1). RNAscope probes were obtained from Advanced Cell Diagnostics (Newark, CA) and used with sections of cochleae from CBA/J wild type mice at P30. Adult cochleae were dissected from the head and fixed overnight at 44°C in 4% PFA in 1x PBS. Fixed adult mouse inner ears were decalcified in 150 mM EDTA for 5-7 days, transferred to 30% sucrose, and then embedded and frozen in SCEM tissue embedding medium (Section-Lab Co, Ltd.). Adhesive film (Section-Lab Co, Ltd.; Hiroshima, Japan) was fastened to the cut surface of the sample in order to support the section and cut slowly with a blade to obtain thin midmodiolar sections. The adhesive film with section attached was submerged in 100% EtOH for 60 seconds, then transferred to distilled water. The adhesive film consists of a thin plastic film and an adhesive and it prevents specimen shrinkage and detachment. This methodology allows for high quality anatomic preservation of the specimen. Frozen tissues were sectioned (10 µm thickness) with a CM3050S cryostat microtome (Leica, Vienna, Austria). Sections were mounted with SCMM mounting media (Section-Lab Co, Ltd., Hiroshima, Japan) and imaged using a 1.4 N.A. objective. Probe information is detailed in both Table 1 and Supplemental Table S1.

#### Immunohistochemistry

For immunohistochemistry of cochlear sections, fixed adult mouse inner ears were decalcified in 150 mM EDTA for 5-7 days, transferred to 30% sucrose, and then embedded and frozen in SCEM tissue embedding medium (Section-Lab Co, Ltd.; Hiroshima, Japan). Adhesive film (Section-Lab Co, Ltd.; Hiroshima, Japan) was fastened to the cut surface of the sample in order to support the section and cut slowly with a blade to obtain 10 µm thickness sections. The adhesive film with sections attached was submerged in 100% EtOH for 60 seconds, then transferred to distilled water. The adhesive film consists of a thin plastic film and an adhesive, which prevents specimen shrinkage and detachment. This methodology allows for high quality anatomic preservation of the specimen along with allowing sectioning at reduced thickness of 0.5 µm. Mid-modiolar sections were obtained from each cochlea where an endocochlear potential recording had been performed.

Fluorescence immunohistochemistry for known stria vascularis cell-type markers was performed as follows. Mid-modiolar sections were washed in PBS then permeabilized and blocked for 1 hour at room temperature in PBS with 0.2% Triton X-100 (PBS-T) with 10% fetal bovine serum (Catalog # A3840001, ThermoFisher Scientific, Waltham, MA). Samples were then incubated in the appropriate primary antibodies in PBS-T with 10% fetal bovine serum, followed by three rinses in PBS-T and labelling with AlexaFluor-conjugated secondary antibodies (1:250, Life Technologies) in PBS-T for 1 hour at room temperature. Where indicated, 4,6-diamidino-2-phenylindole (1:10,000, Life Technologies) was included with the secondary antibodies to detect nuclei. Organs were rinse in PBS three times and mounted in SlowFade (Invitrogen). Specimens were imaged using a Zeiss LSM710 confocal microscope. Sections were mounted with SCEM mounting medium (Section-Lab Co, Ltd., Hiroshima, Japan). Primary antibodies used included rabbit anti-KCNJ10 (Alomone Labs, Cat# APC-035, polyclonal, dilution 1:200), rabbit anti-CLDN11 (Life Technologies, Cat# 364500, polyclonal, dilution 1:200), goat anti-SLC12A2 (Santa Cruz Biotech, Cat# sc-21545, polyclonal, dilution 1:200), goat anti-KCNQ1 (Santa Cruz Biotech, Cat# sc-10646, polyclonal, dilution 1:200), Phalloidin AlexaFluor 647 (Invitrogen, Cat# A22287, dilution 1:250).

## Supporting information

Supplemental Results

## Statistical analysis

Statistical analysis for single-cell RNA-Seq is described in the detailed methods.

## Data and Software Availability

All data generated in these studies have been deposited in the Gene Expression Omnibus (GEO) database (GEO Accession ID: GSE136196) and will be available upon publication. We are also in the process of uploading the data into the gene Expression Analysis Resource (gEAR), a website for visualization and comparative analysis of multi-omic data, with an emphasis on hearing research (https://umgear.org).

## Acknowledgements

This research was supported (in part) by the Intramural Research Program of the NIH, NIDCD to M.H. (DC000088) and R.J.M. (DC000086). This research was made possible through the NIH Medical Research Scholars Program, a public-private partnership supported jointly by the NIH and contributions to the Foundation for the NIH from the Doris Duke Charitable Foundation (DDCF Grant #2014194), the American Association for Dental Research, the Colgate-Palmolive Company, Genentech, Elsevier, and other private donors. The authors would like to acknowledge Thomas B. Friedman, Matthew W. Kelley and Lisa Cunningham who provided helpful feedback and review of this paper. The authors acknowledge Alan Hoofring for his illustrations. This study utilized the high-performance computational capabilities of the Biowulf Linux cluster at the National Institutes of Health, Bethesda, MD. (http://biowulf.nih.gov)

## Ethics Statement

All animal experiments and procedures were performed according to protocols approved by the Animal Care and Use Committee of the National Institute of Neurological Diseases and Stroke and the National Institute on Deafness and Other Communication Disorders, National Institutes of Health.

## Supplemental Figures

**Supplemental Figure S1.**
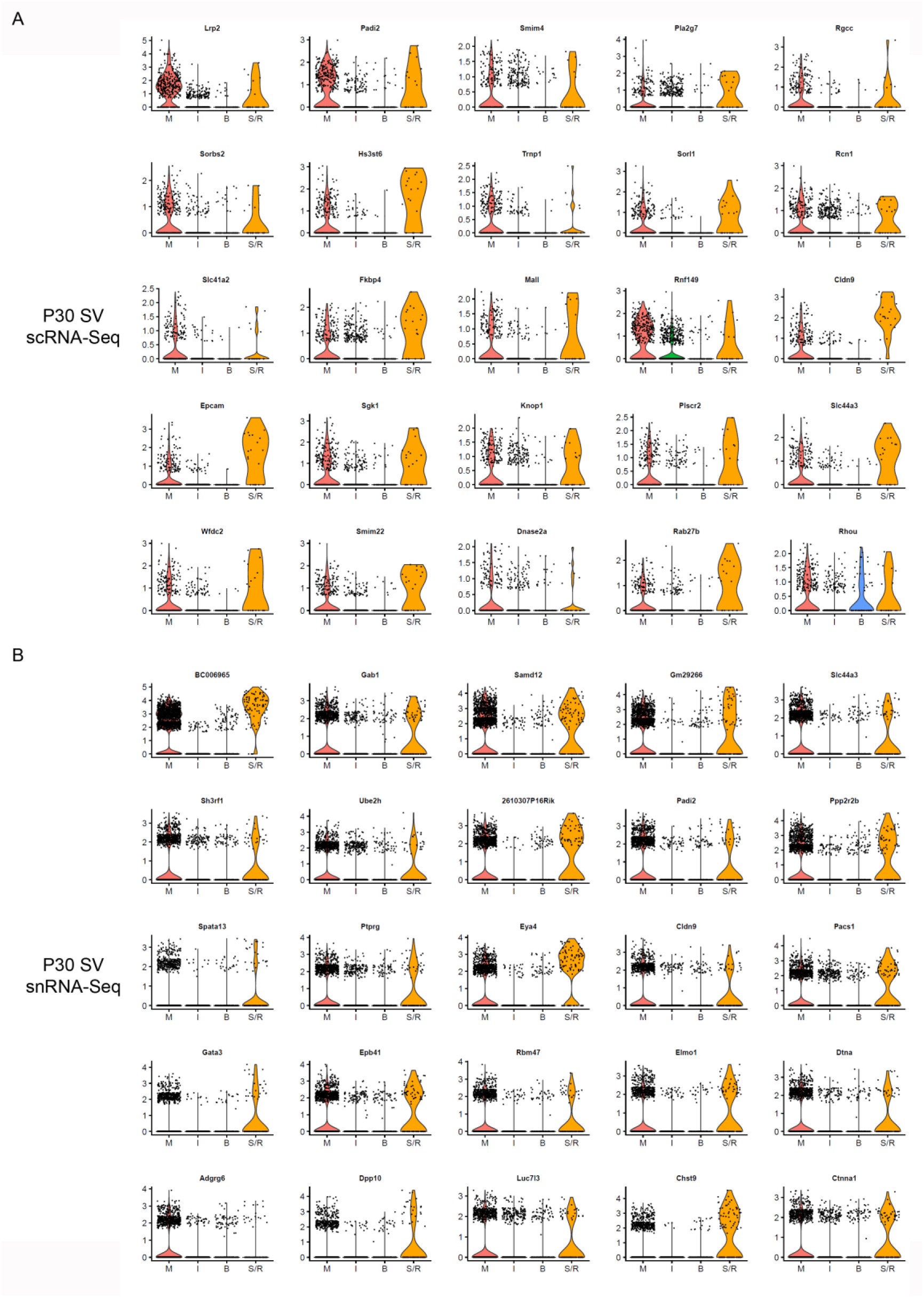
Shared gene expression between marginal and spindle cells. (A), Candidate genes identified in the scRNA-Seq dataset expressed by marginal (M) and spindle / root (S/R) cells. (B), Candidate genes identified in the snRNA-Seq dataset expressed by marginal (M) and spindle / root (S/R) cells. Intermediate cells (I), basal cells Violin plots are displayed with normalized counts on the vertical axis and cell types arrayed along the horizontal axis.

**Supplemental Figure S2.**
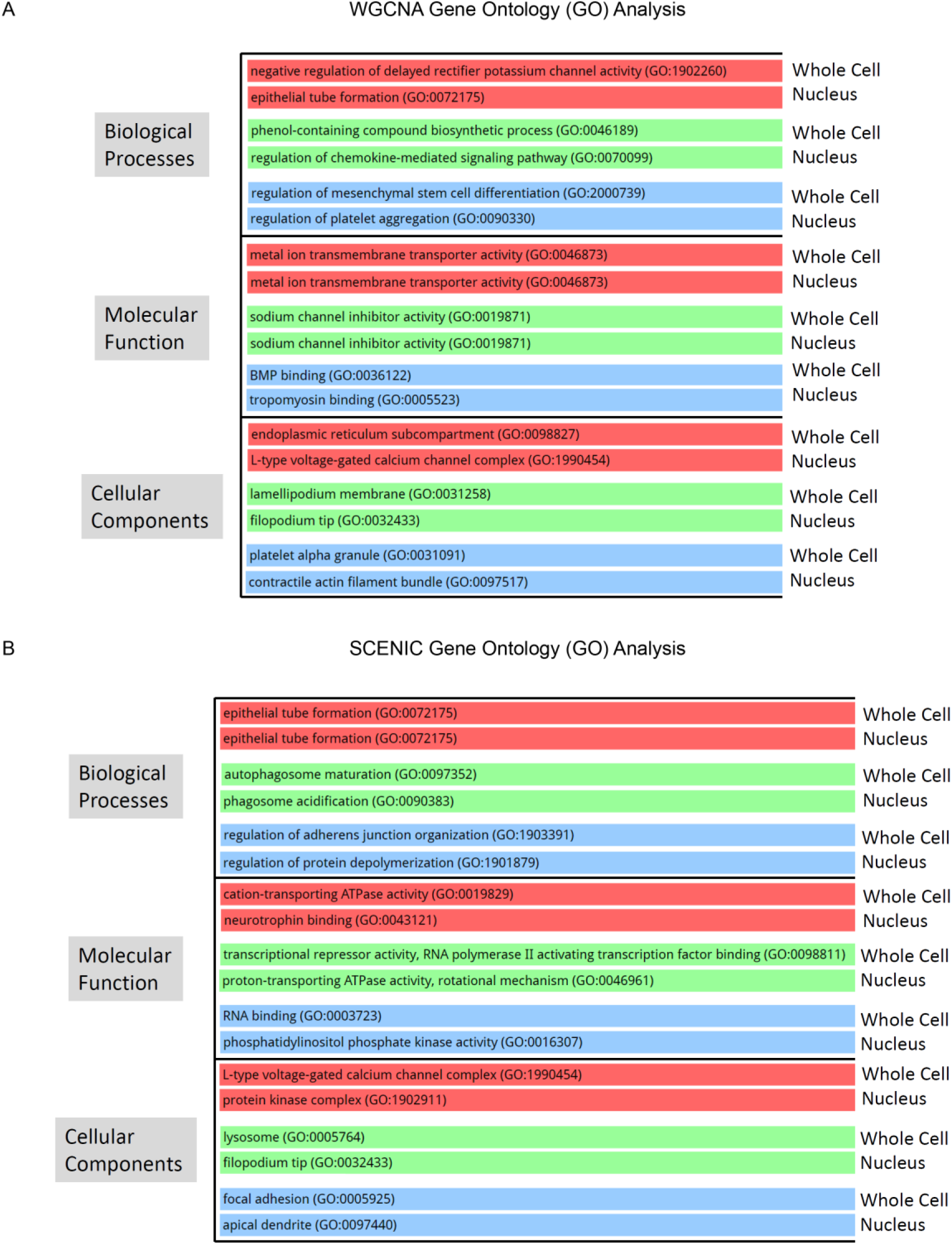
Top gene ontology (GO) terms identified by GO analysis with Enrichr in cell type-specific WGCNA modules and cell type-specific SCENIC regulons. (A), GO biological process, molecular function, and cellular component analysis of WGCNA modules from both the scRNA-Seq and snRNA-Seq datasets reveal gene set enrichment in marginal cells (red), intermediate cells (green), and basal cells (blue). (B) GO biological process, molecular function, and cellular component analysis of SCENIC regulons from both the scRNA-Seq and snRNA-Seq datasets reveal gene set enrichment in marginal cells (red), intermediate cells (green), and basal cells (blue).

**Supplemental Figure S3.**
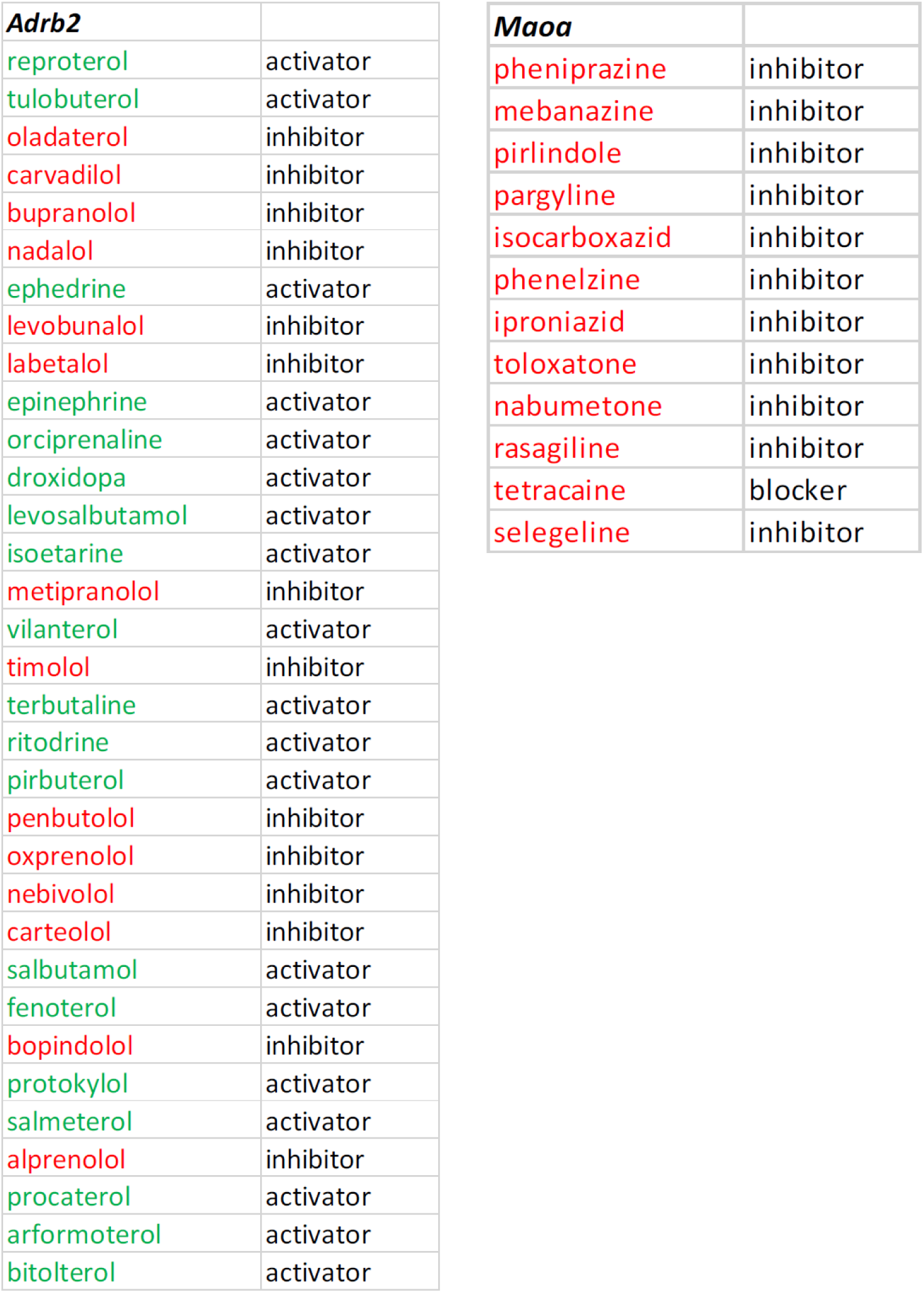
Pharos-identified drugs with Adrb2 and Maoa gene activity. Drugs are displayed along with their mechanism of action. Activators are in green and inhibitors are in red.

## Supplemental Tables

**Supplemental Table S1.** RNAScope *in situ* probes

| Probe | Company (Region targeted) | Catalog # | Accession No. |
| --- | --- | --- | --- |
| *Abcg1* | Advanced Cell Diagnostics (3680 – 4595) | 422221-c2 | NM_009593.2 |
| *Atp13a5* | Advanced Cell Diagnostics (2060 – 4007) | 417211 | NM_175650.4 |
| *Atp1b2* | Advanced Cell Diagnostics (968 – 2027) | 417131 | NM_013415.5 |
| *Bmyc* | Advanced Cell Diagnostics (6 – 874) | 593551-C3 | NM_023326.2 |
| *Cd44* | Advanced Cell Diagnostics (2 – 947) | 479191 | NM_009851.2 |
| *Esrrb* | Advanced Cell Diagnostics (437 – 1577) | 565951-c3 | NM_001159500.1 |
| *Heyl* | Advanced Cell Diagnostics (425 – 1345) | 446881 | NM_013905.3 |
| *Kcne1* | Advanced Cell Diagnostics (6 – 929) | 541301 | NM_008424.3 |
| *Kcnj13* | Advanced Cell Diagnostics (2 – 1098) | 412551-c3 | NM_001159500.1 |
| *Kcnj16* | Advanced Cell Diagnostics (2 – 961) | 492481-c3 | NM_001252207.1 |
| *Kcnq1* | Advanced Cell Diagnostics (6 – 874) | 420481-c2 | NM_008434.2 |
| *Lrp2* | Advanced Cell Diagnostics (1230 – 2350) | 487751 | NM_001247984.1 |
| *Met* | Advanced Cell Diagnostics (3341 – 4309) | 405301-c2 | NM_008591.2 |
| *Nr2f2* | Advanced Cell Diagnostics (1532 – 3193) | 480301 | NM_009697.3 |
| *Nrp2* | Advanced Cell Diagnostics (2001 – 2870) | 500661 | NM_001077406.1 |
| *P2rx2* | Advanced Cell Diagnostics (228 – 1394) | 443681-c2 | NM_153400.4 |
| *Pax3* | Advanced Cell Diagnostics (697 – 1675) | 455801 | NM_008781.4 |
| *Sox8* | Advanced Cell Diagnostics (931 – 1876) | 454781 | NM_011447.3 |
| *Slc26a4* | Advanced Cell Diagnostics (795 – 1776) | 452491 | NM_011867.3 |

