## Supplemental Results for "Single cell and single nucleus RNA-Seq reveal cellular heterogeneity and homeostatic regulatory networks in adult mouse stria vascularis"

### Review of novel SV genes

*Abcg1* is a lipid transporter that appears to be involved in regulation of cholesterol homeostasis at the blood-brain barrier and mutations in Abcg1 have been implicated in impaired monocyte cholesterol clearance in age-related macular degeneration resulting in vision loss (Ban et al., 2018; Kober et al., 2017; Tarling, 2013). Prior to this study, *Abcg1* expression had not been demonstrated in the cochlea. Malgrange and colleagues have suggested the possibility that cholesterol homeostasis plays a role in the development of SNHL and that genes involved may offer the possibility of therapeutic approaches to treat or prevent SNHL (Malgrange, Varela-Nieto, de Medina, & Paillasse, 2015). *Heyl* is a downstream effector of the Notch signaling pathway and has been shown to be expressed in the developing organ of Corti, appearing to be responsible for maintaining the fate of cochlear supporting cells (Doetzlhofer et al., 2009; McGovern, Zhou, Randle, & Cox, 2018). However, prior to this study, *Heyl* expression had not been demonstrated in the adult cochlea, although presumably it plays a role maintaining cell fate. In addition to its role in axon guidance, *Nrp2* has been recently demonstrated to be expressed in the cuboidal cells of the avian tegmentum vasculosum, the avian equivalent to the mammalian SV (Coate, Spita, Zhang, Isgrig, & Kelley, 2015; Scott, Yue, Biesemeier, Lee, & Fekete, 2019). A conditional knockout mouse of an *Nrp2* paralog, *Nrp1*, has been associated with enlarged microvessels of the SV and *Nrp1* is expressed in the developing SV at postnatal day 5 (Salehi et al., 2017). *Kcnj13*, encodes a potassium inwardly-rectifying channel known as Kir7.1, has been identified as being expressed by adult cochlear hair cell and supporting cell transcriptome profiles but has not been demonstrated to be expressed in the adult SV (Liu et al., 2018, 2014). *Sox8* is a transcription factor involved in initial signaling and maintenance of *Sox10* expression in the otic placode during inner ear development (Betancur, Sauka-Spengler, & Bronner, 2011). While *Sox8* has been shown to be expressed in the developing chicken otocyst in the region of the tegmentum vasculosum (Sinkkonen et al., 2011), the equivalent to the mammalian SV, it has not been previously demonstrated in the mammalian stria vascularis, much less the adult SV. *Nr2f2* is an orphan steroid/thyroid hormone nuclear receptor with an essential role in angiogenesis during development and known expression in embryonic and adult mouse cochlear and vestibular sensory epithelia (Tornari, Towers, Gale, & Dawson, 2014). Expression of *Nr2f2* has not been previously demonstrated in the SV. *Kcnj16*, encodes a potassium inwardly-rectifyng channel known as Kir5.1, has been identified as being expressed by adult cochlear hair cell and supporting cell transcriptome profiles but has not been demonstrated to be expressed in the adult SV (Liu et al., 2018, 2014). *P2rx2*, encodes the P2X2 ATP-sensitive non-specific cation channel which is known to be expressed in outer sulcus cells but previously described as being absent in the SV (Järlebark, Housley, Raybould, Vlajkovic, & Thorne, 2002; Järlebark, Housley, & Thorne, 2000). We demonstrate *P2rx2* transcript expression in SV spindle cells (Figure 3E, 3F). Finally, *Atp13a5* is a P5 ATPase that has been demonstrated to be highly expressed in the adult mouse brain (Schultheis et al., 2004; Weingarten, Dave, Li, & Crawford, 2012). *Atp13a5* has been shown to be expressed in transcriptome profiles from whole mouse cochlea from early postnatal mice but has not been previously localized within the SV (Son et al., 2012).
